# Regioselective control of biocatalytic C–H activation and halogenation

**DOI:** 10.1101/2022.08.04.502814

**Authors:** Elijah N. Kissman, Monica E. Neugebauer, Kiera H. Sumida, Cameron V. Swenson, Nicholas A. Sambold, Jorge A. Marchand, Douglas C. Millar, Michelle C.Y. Chang

## Abstract

Biocatalytic C–H activation has the potential to merge enzymatic and synthetic strategies for bond formation. Fe^II^/αKG-dependent halogenases are particularly distinguished for their ability both to control selective C-H activation as well as to direct group transfer of a bound anion along a reaction axis separate from oxygen rebound, enabling the development of new transformations. In this context, we elucidate the basis for selectivity of enzymes that perform selective halogenation to yield 4-Cl-lysine (BesD), 5-Cl-lysine (HalB), and 4-Cl-ornithine (HalD), allowing us to probe how regioselectivity and chain length selectivity are achieved. We now report the crystal structure of the HalB and HalD, revealing the key role of the substrate-lid in positioning the substrate for C_4_ vs C_5_ chlorination and recognition of lysine vs ornithine. Targeted engineering of the substrate-binding lid further demonstrates that these selectivities can be altered or switched, showcasing the potential to develop halogenases for biocatalytic applications.

## INTRODUCTION

The use of enzymes as biocatalysts has expanded the scope of chemical synthesis by introducing new avenues for reducing synthetic steps to minimize waste and cost for industrial synthesis^1–5^. Given their high selectivity and compatibility with mild aqueous conditions compared to chemical catalysts, the integration of enzymes with traditional synthetic chemistry has changed the landscape of what can be achieved by telescoped and single-pot synthetic routes^6–8^. While several classes of enzymes, such as ketoreductases, transaminases, hydrolases, and lyases, can be reliably incorporated into commercial synthesis, there remains a need for new reaction classes in order to blend biological and chemical catalysis^9–14^. In this regard, selective C-H activation and functionalization could enable new C-C bond forming connections to rapidly assemble complex molecules by interfacing enzymatic selectivity for C-H activation with the breadth of downstream synthetic reactions used for bond making and cross coupling (**Fig. 1A**).

**Figure 1.**
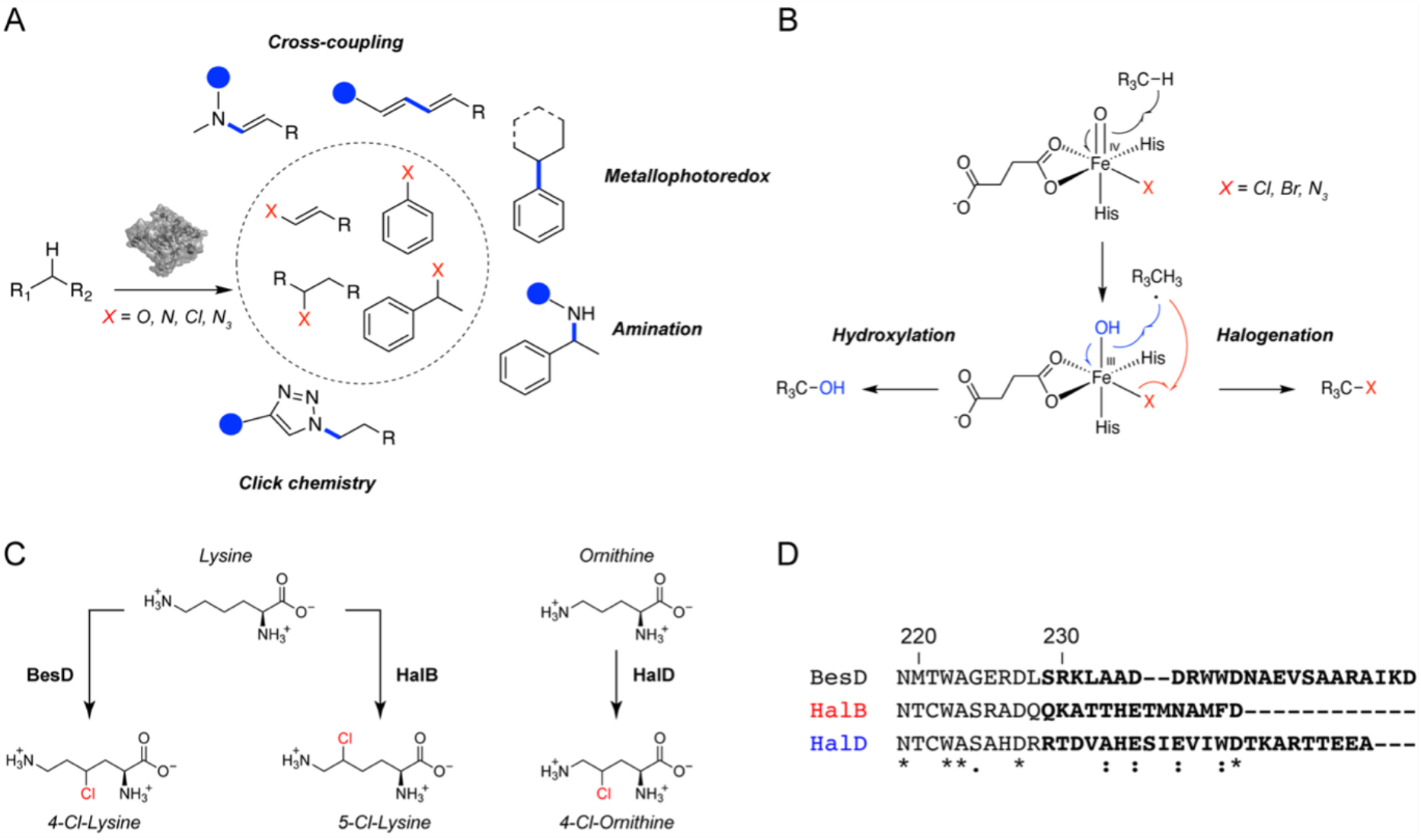
Radical amino acid halogenase selectivity. **(A)** Enzymatic C-H activation has been utilized for selective C-C and C-heteroatom bond formation. These products have been combined with downstream synthetic and enzymatic reactions to form pharmaceutically relevant intermediates. **(B)** Control of the reactive Fe^IV^-oxo intermediate can enable the delivery of alternate anions such as azide. Engineering the enzymes while avoiding off-pathway hydroxylation has been an ongoing challenge. **(C)** Regioselectivity of lysine halogenases. BesD chlorinates lysine at C_4_ to yield 4-Cl-lysine. HalB chlorinates lysine at C_5_ to yield 5-Cl-lysine. HalD displays substrate selectivity for ornithine, which is one carbon shorter than lysine **(D)** The C-terminus of the halogenases vary significantly depending on the substrate and regioselectivity. HalB and HalD align more closely while BesD is longer and shares fewer similar residues. The annotated C-terminal region is shown in bold, and residue numbering is based on the sequence of BesD.

Although C-H activation reactions have found high utility in synthetic chemistry, site selectivity can be challenging to achieve as the weakest or most sterically-accessible C-H bond is typically activated by the catalyst, it can be difficult to prevent undesired activation of C-H bonds of similar strength, and functional group compatibility can be low. To address these challenges, chemical catalysts have been designed to use ligand architecture or spatial orientation to achieve selectivity^15–23^. In contrast, enzymes routinely achieve both stereo- and regioselectivity when presented with similar sites of equivalent reactivity since their three-dimensional chiral architecture naturally creates asymmetry. Although the most common outcome for enzymatic C-H activation is O atom transfer, enzymes have been harnessed for myriad useful reactions including C–C bond formation,^24,25^ amination,^26^ azidation^27^, silation,^28^ and borylation.^29^ Thus, C-H activating enzymes provide a promising starting point for diversifying and expanding the role of biocatalysis in chemistry.

Many classes of metalloenzymes, including cytochrome P450s^10,12,30,31^, Fe/αKG-dependent enzymes ^12,32–38^, and Rieske enzymes^12,39–41^ have been used for biocatalytic C-H activation reactions via H atom abstraction to form a substrate radical intermediate. Radical-based transformations have particularly stringent requirements for active-site geometry; not only is there a strong distance-dependence for hydrogen atom abstraction, but the reactive nature of the intermediates can lead to a wide range of side reactions if not precisely coordinated^42^. We are particularly interested in studying Fe/αKG-dependent halogenases as a paradigm for understanding the geometric active-site constraints that control site and reaction selectivity for C-H activation and rebound. In Fe/αKG-dependent halogenases, substrate radical formation by a ferryl intermediate and subsequent substrate radical functionalization with halide (X = Cl, Br) occur along different vectorial paths (**Fig. 1B**)^43–47^. This behavior leads to flexibility in transferring new and abiotic anions, such as azide^27^, but also presents challenges for active site engineering.

Biocatalytic applications of Fe/αKG-dependent halogenases have been limited by the relatively small number of halogenases discovered to date as well as the difficulty in engineering their substrate selectivity without altering their reaction pathway towards hydroxylation. The first reported radical halogenase family was found to require a carrier-protein tethered substrate^48–55^. However new families of radical halogenases have been discovered to act on freestanding substrates, including alkaloids^56^, nucleotides^57^, and amino acids.^58^ These families collectively provide promising new scaffolds for diversification in both substrate and reaction range.^37,59^ We discovered a new family of Fe^II^/αKG-dependent halogenases (BesD) that carries out the regio- and stereoselective C(*sp*^3^)-H activation of simple amino acid substrates without the need for a carrier protein.^60^ Given the privileged role of amino acids as building blocks for biosynthesis, catalyst design, and drug discovery^61^, elucidation of the sequence and structural determinants that govern the substrate selectivity of BesD halogenases can allow us to tap their potential use in biocatalytic applications.

Toward this goal, we investigate enzymes within the BesD amino acid halogenase family that perform subtly different reactions to selectively yield either 4-chlorolysine (BesD), 5-chlorolysine (HalB), or 4-chloroornithine (HalD) (**Fig. 1C**). Through bioinformatics analysis and X-ray crystallography of HalB, we identify a C-terminal substrate-binding lid that undergoes an unprecedented conformational change to control regioselectivity and substrate selectivity within this family. To understand how BesD and HalB chlorinate lysine at C_4_ and C_5_, respectively, we solve the lysine-bound crystal structure of HalB for comparison to BesD^60^, which reveals how structurally distinct substrate-binding lids tune the substrate conformation within the active site to achieve divergent regioselective outcomes. Through site-directed mutagenesis and fluorogenic screening, we identify mutations that disrupt regioselectivity and support the proposed role of the substrate-binding lid in controlling reactivity. Next, we solve the crystal structure of the ornithine halogenase, HalD, which has a 40-fold lower K_M_ for the 5-carbon substrate, ornithine, compared to the 6-carbon substrate, lysine. The ornithine and lysine halogenases are structurally similar except at the lid domain, which closes over the substrate in the active site. Structure-guided engineering of the HalD lid expands the substrate binding pocket, enabling HalD to chlorinate lysine with a K_M_ comparable to that of HalB. Taken together, this work provides fundamental insights into the basis of regioselective halogenation of unactivated C_sp3_–H bonds and suggests that the substrate binding lid may be a promising engineering target for future efforts to tailor radical halogenases for the biocatalytic chlorination of a broad range of substrates.

## RESULTS

### Amino acid halogenases have variable C-termini which are involved in substrate binding

Given the synthetic utility of reactions catalyzed by Fe/αKG-dependent enzymes and halogenation in particular, a detailed understanding of how these enzymes recognize small molecule substrates is key to expanding their biocatalytic applications. Indeed, the substrate selectivities found within the BesD radical halogenase family allow us explore the molecular basis for how regioselectivity and chain length selectivity is controlled (**Fig. 1C**)^60^. We previously solved the crystal structure of the lysine 4-chlorinase, BesD, which shows that the C-terminal domain (residues 229–252) forms a lid over the substrate in the active site.^60^ This closed active site appears to be conserved in other structures of Fe/αKG-dependent enzymes and in some other radical-utilizing enzymes that accept small molecule rather than protein-bound substrates^62^. Interestingly, a sequence alignment of BesD (4-chlorolysine), HalB (5-chlorolysine), and HalD (4-chloroornithine), revealed that the active-site residues are highly conserved except those within the C-terminal lid, which varies in length and sequence (**Fig. 1D, Supplementary Fig. 1**). Given the role of the C-terminal domain in forming the active site and its variability, we decided to further explore its role in substrate selectivity.

Towards this end, we initiated structural studies of HalB, which chlorinates the same lysine substrate as BesD but at C_5_ instead of C_4_.^60^ Notably, the HalB C-terminus (15 amino acids) is 40% shorter than that of BesD (25 amino acids) and the two C-terminal sequences align poorly. HalB crystals formed in succinate-phosphate-glycine (SPG) buffer, diffracted to 2.05 Å, and contained two copies of HalB per asymmetric unit (**Supplementary Table 2**). Strikingly, the two copies of HalB adopted surprisingly different conformations depending on the presence or absence of a bound amino acid. In the apo chain A, no amino acid is found in the active site and the C-terminal domain (residues 238–251) that forms the substrate lid is disordered (**Fig. 2C**). In chain B, density for glycine is present in the enzyme active site in the position expected for the substrate lysine, likely due to the high concentration of glycine in the crystallization conditions (21.8 mM) (**Fig. 2A, B**). In the presence of an amino acid, the C-terminal domain becomes structured. In the absence of an amino acid, the antiparallel beta strands (residues 68–88) that bind the carboxylate of the substrate via Arg83 are instead projected away from the active site to adopt an extended alpha helical structure. Although substrate-covering lids have been identified in other Fe/αKG-dependent enzymes^45,62^, the observed helix-to-sheet transition is unique to HalB. This snapshot of HalB in two different states suggests that HalB undergoes a conformational change upon substrate binding with the C-terminal lid forming part of the active site to participate in substrate recognition and catalysis.

**Figure 2.**
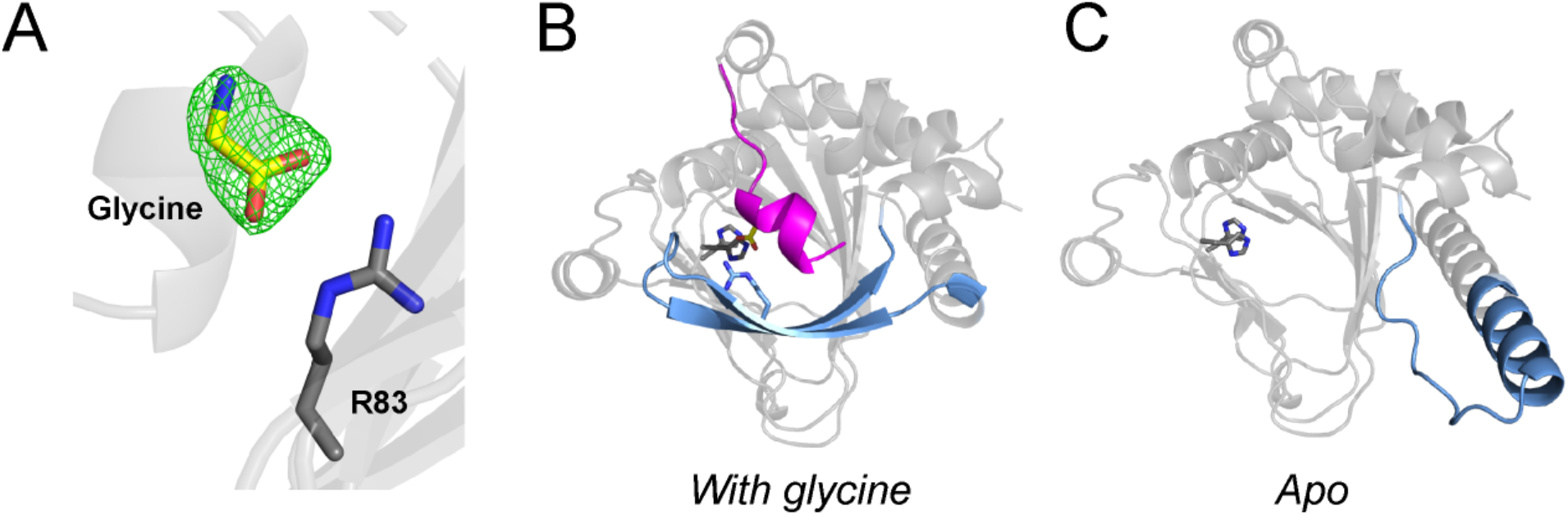
HalB undergoes a conformational change upon substrate binding. **(A)** Crystal structure of glycine-bound Chain A of HalB (2.05 Å) with F_o_-F_c_ omit map (green mesh contoured at 1.5 σ) for glycine (yellow). **(B)** Structure of glycine-bound HalB with the C-terminal domain (residues 238–251) shown in magenta and the antiparallel beta strands containing Arg83 shown in blue (residues 68–88). **(C)** Structure of apo chain A of HalB with the alpha helical domain in blue. The residues in the alpha helical domain correspond to the residues in the beta strands in **(B)**. The C-terminal domain (residues 238–251) is disordered and therefore not modelled.

### Lysine-bound crystal structure of HalB

To investigate how the C-terminal lid of HalB interacts with its native substrate, we next solved the lysine-bound structure of HalB to 1.90 Å by screening for crystal formation in conditions containing lysine instead of glycine (**Supplementary Table 3**). Under the new crystallization conditions, density for the lysine ligand is clearly visible in the active site of HalB (**Fig. 3A**). The structures of HalB and BesD are similar except at their C-terminal lids, which close over the substrate in the active site (**Fig. 3B**). While the C-terminus of BesD is longer and lacks secondary structure, the C-terminus of HalB is shorter and α-helical. An overlay of the crystal structures of HalB and BesD reveals that the enzymes have conserved carboxylate and α-amine binding residues (**Fig. 3C**). As a result, the lysine substrate is located in a similar position in the active site and is not translationally shifted in HalB compared to BesD. However, we observed striking differences in the conformation of lysine within the active sites. In BesD, C_4_ of lysine is oriented toward the Fe complex. In contrast, C_5_ of lysine is instead presented toward the complex in HalB, while C_4_ is pointed away. The conformations of lysine within each structure are consistent with the observed regioselective outcomes for the two enzymes (**Fig. 2C**).

**Figure 3.**
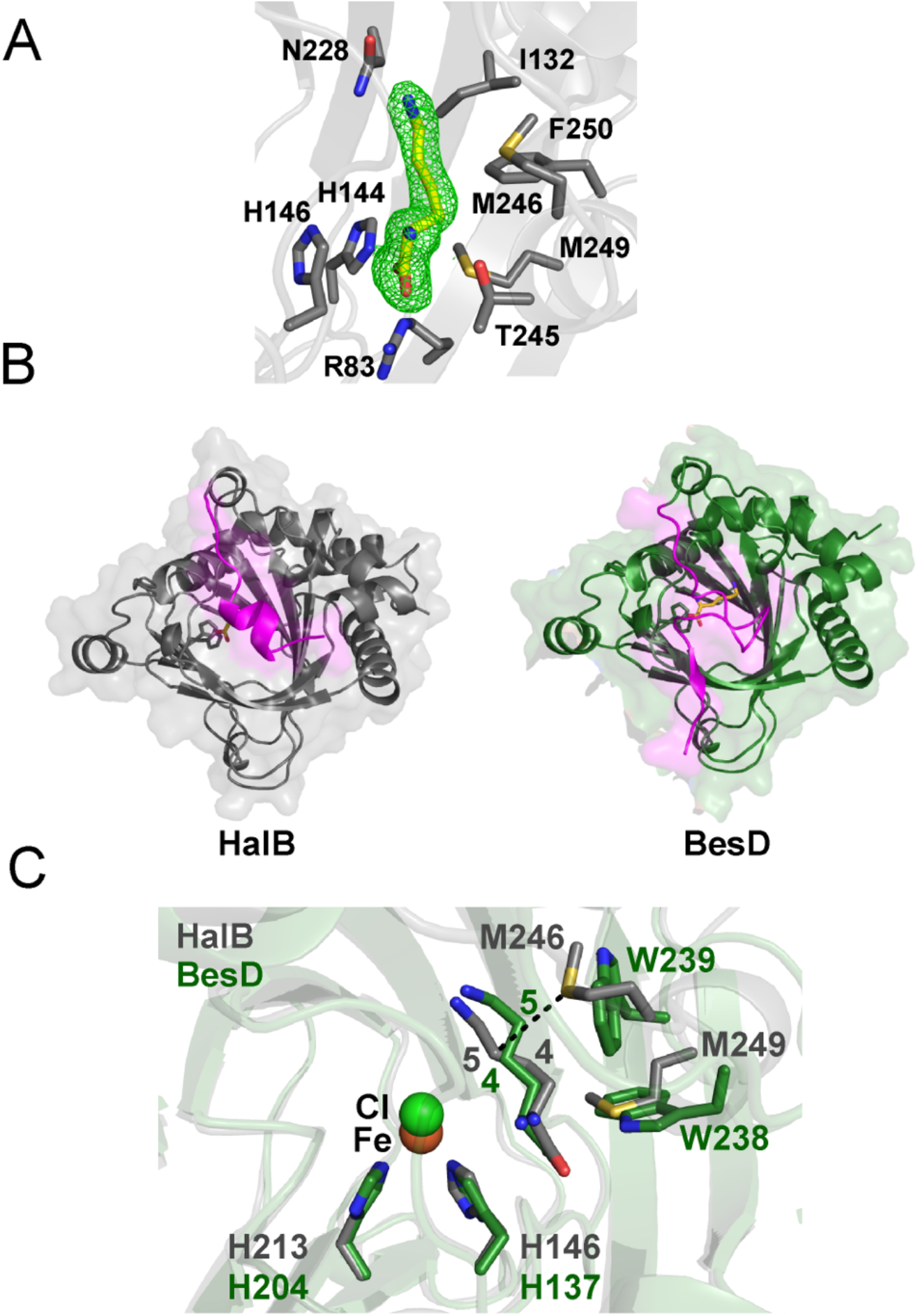
Structural basis of regioselectivity. **(A)** Crystal structure of HalB (1.9 Å) with F_o_-F_c_ omit map (green mesh, contoured at 1 σ) for lysine (yellow). Active site residues of HalB are shown as grey sticks. **(B)** Comparison of the overall structures of HalB (1.90 Å, grey) and BesD (1.95 Å, green, PDB 6NIE). The enzymes share high structural similarity except at the substrate covering lid (magenta). **(C)** Structural alignment of HalB (grey) and BesD (green). Although the carboxylate and amine of lysine are in a similar position in both cases, the sidechain samples a different conformation in the two enzymes. In BesD, C_4_ is oriented toward chloride in the active site, while C_5_ is oriented away. In contrast, HalB orients C_5_ toward chloride. The substrate conformation is influenced by residues in the substrate covering lids of BesD and HalB. In particular, Met246 in HalB makes a steric contribution to the reaction outcome.

Notably, the key residues that contribute to the conformational differences of lysine are located on the C-terminal substrate-binding lids. In BesD, Trp239 stacks against the aliphatic side chain of lysine in the active site. In HalB, this Trp residue is absent. Instead, Met246 protrudes into the active site and presents C_5_ of lysine toward the Fe complex, making it accessible for halogenation. Thus, Met246 makes a steric contribution to halogenation of lysine at C_5_. This mechanism is distinct from some other amino-acid modifying enzymes which enforce regioselectivity by translationally shifting the substrate. For example, lysine 3-hydroxylase KDO1 binds the substrate carboxylate with Arg332 while lysine 4-hydroxylase KDO5 pulls the substrate farther into the active site to interact with Arg145 which is absent in KDO1.^62^ In comparison, while lysine adopts an altered conformation in the active site of BesD and HalB (all-atom RMSD 1.112 Å), there is no significant shifting of the key binding determinants, with the α-carboxylate, α-amine, and ε-amine shifted by just 0.2, 0.4, and 0.6 Å respectively.

In addition to substrate selectivity controlled by hydrogen atom abstraction, radical halogenases also need to solve the challenge of rebound selectivity in the next step with the halide ligand over the thermodynamically preferred pathway of hydroxyl ligand rebound (**Supplementary Fig. 2**).^36,63^ The selectivity of rebound has been hypothesized to be derived from precise control over the substrate radical positioning relative to the two potential rebounding species to allow for the more difficult halogenation reaction to occur.^43–46,64,65^. Although pairs of Fe^II^/αKG-dependent amino acid hydroxylases with regio-divergent outcomes have been reported^62,66–68^, it is remarkable that the two halogenases, BesD and HalB, overcome the additional challenging of precisely orienting lysine within the active site to achieve regioselectivity without compromising selectivity for halogenation over hydroxylation.

### Investigating the role of Met246

To probe the significance of Met246 on regioselectivity in HalB, we generated a M246 NNK library to determine whether mutagenesis could perturb the regioselectivity of HalB for modifying C_5_ and result in chlorination of C_4_ as well. To assay the variants, we employed a fluorogenic screen^69^ based on the role of 4-Cl-lysine as an intermediate in the biosynthesis of propargylglycine (Pra), a terminal-alkyne containing amino acid (**Fig. 4A**)^58^. In this pathway, 4-Cl-lysine produced by BesD is a substrate for the downstream enzymes BesC and BesB, which form Pra^58^. If lysine is chlorinated at C_5_ instead, Pra is not formed, since 5-Cl-lysine is not a substrate for BesC and BesB. As a result, only cells that produce 4-Cl-lysine enable copper-catalyzed azide-alkyne cycloaddition (CuAAC) with the fluorogenic probe CalFluor 488 (**Supplementary Fig. 3**)^70^. This screen can therefore be used to determine whether mutagenesis of Met246 in HalB disrupts regioselective halogenation of C_5_.

**Figure 4.**
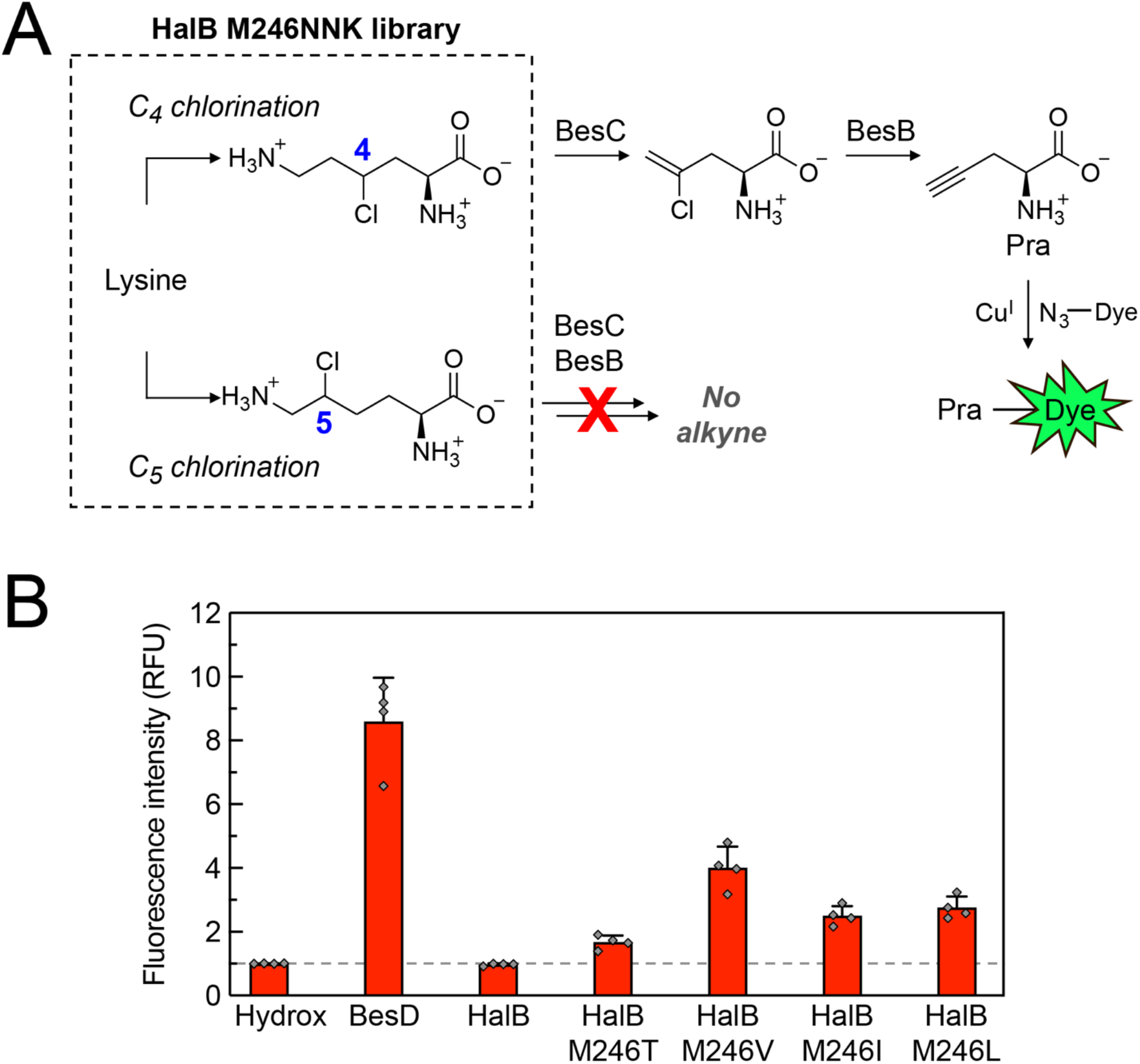
Screening the HalB library for perturbation in regioselectivity. **(A)** Strategy for screening the M246NNK library for variants that perform halogenation. In two enzymatic steps, 4-Cl-lysine can be converted into Pra, which contains an alkyne functional group that can be leveraged for CuAAC chemistry with a fluorogenic probe. Variants that only perform chlorination of C_5_ yield no fluorescence, as 5-Cl-lysine is not a substrate for the enzymes in the alkyne biosynthesis pathway. **(B)** Cells expressing a lysine 4-hydroxylase (Hydrox), lysine 4-halogenase (BesD), lysine 5-halogenase (HalB), or a variant of HalB, along with downstream enzymes BesB and BesC were grown in 96-well plates at 16°C for 48 hours, at which time the cultures were supplemented with lysine and αKG. After an additional 2 d of growth, cells were pelleted by centrifugation and the supernatant was analyzed for Pra by reaction with CalFluor 488, BTTAA, sodium ascorbate, and copper (II) sulfate. Following incubation in the dark for 15 min, fluorescence intensity was measured at room temperature (λ_ex_ = 485 nm, λ_em_ = 528 nm). Mean and error are shown for n = 4 technical replicates.

We transformed the HalB M246 NNK library into cells containing *besC* and *besB* and inoculated 168 colonies into a 96-well plate. The library was screened for alkyne formation alongside wild type BesD, HalB, and Hydrox, a negative control which makes 4-hydroxylysine^4^. After four days of expression, the cells were removed by centrifugation, and the supernatants were analyzed by CuAAC with CalFluor 488 (**Supplementary Fig. 4**). As expected, cells expressing HalB (lysine 5-halogenase) yielded no fluorescence increase relative to the negative control, consistent with the lack of 4-Cl-lysine production. In contrast, cells expressing BesD (lysine 4-halogenase) yielded an 8-fold increase in fluorescence. The screen identified 26 variants among the NNK library that yielded a 2-fold increase in fluorescence intensity relative to Hydrox (**Supplementary Fig. 4**). Plasmids from these alkyne-producing cells were isolated and sequenced to identifying mutations that enabled 4-Cl-lysine formation. M246L, M246I, M246V, and M246T appeared among the 26 sequenced hits (**Supplementary Table 4**). The results suggest that replacement of Met246 with smaller and/or nonpolar residues may alleviate steric constraints that position C_5_ close to the catalytic Fe, thereby enabling chlorination of C_4_ as well.

### Characterization of HalB variants

We repeated the assay with replicates of each of these clones and determined that M246V, M246L, M246I, and M246T yielded a 4-fold, 2.75-fold, 2.5-fold, and 1.67-fold fluorescence increase, respectively, relative to HalB (**Fig. 4B**). We selected the HalB mutants with the greatest signal from the click screen (HalB M246V, HalB M246L, and HalB M246I) for further characterization. We first evaluated the extent to which the mutations perturbed regioselectivity by comparing the amount of Pra produced in each condition (**Supplementary Fig. 5**). The lysine 4-halogenase, BesD, produced 6.1 ± 0.7 μM Pra in the spent media. HalB M246V resulted in the greatest Pra production, yielding 1.5 ± 0.4 μM Pra. The M246I and M246L mutants produced 0.7 ± 0.1 μM and 0.5 ± 0.1 μM Pra, respectively. The data suggest that M246 is important for regioselectivity and that mutagenesis of this residue leads to aberrant activation of C_4_, albeit at levels much lower than observed in the native lysine 4-halogenase.

**Figure 5.**
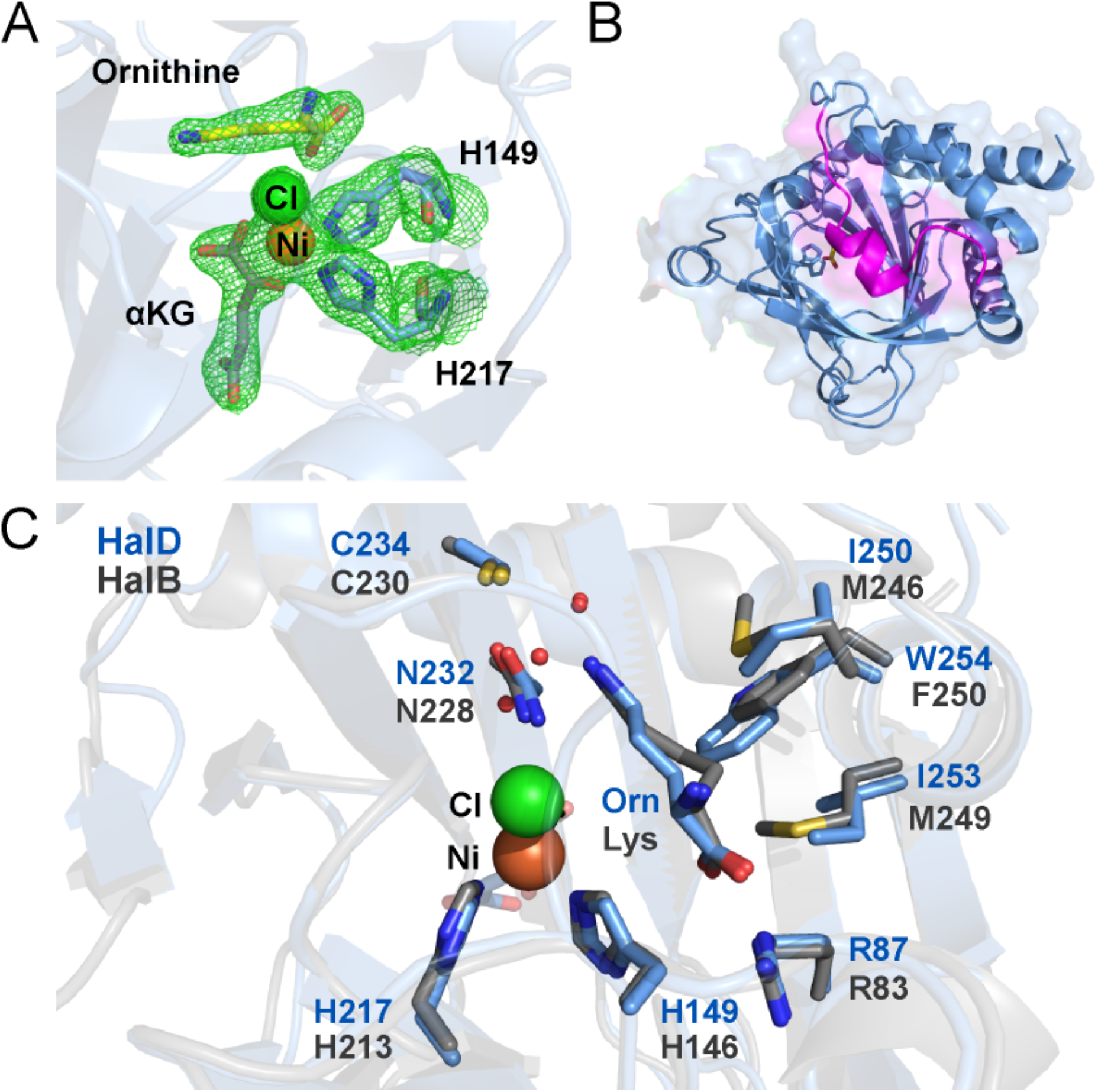
Structural basis of chain length selectivity. **(A)** Density for ornithine in the active site of HalD (2.0 Å). F_o_-F_c_ omit map (green mesh, contoured at 1.5 σ) for ornithine. (B) Overall fold of the ornithine halogenase, HalD. The substrate covering lid is shown in magenta. (C) Structural alignment of HalB (1.9 Å, grey) and HalD (2.0 Å, blue). In contrast to lysine in HalB, the ornithine substrate of HalD is present in an extended conformation, with C_4_ poised for chlorination. HalD is unable to accommodate lysine in the active site because of steric strain imposed primarily by the substitution of Trp254 in place of phenylalanine.

We next investigated whether the mutations that enabled C_4_ chlorination might also lead to diminished overall activity. We purified HalB M246V, HalB M246L, and HalB M246I and performed steady-state kinetic analysis using an assay that couples succinate formation by the halogenase to NADH oxidation^71^. We found that all three mutants had a decreased *k*_*cat*_ and *K*_*M*_ relative to the wild type HalB enzyme, consistent with a perturbation of lysine binding in the enzyme active site (**Supplementary Fig. 6**). These kinetic data were consistent with mass spectrometry data showing that the M246 mutants exhibited as much as a two-fold decrease in production of 5-chlorolysine compared to HalB (**Supplementary Fig. 7**). The promiscuity of the HalB variants supports the significance of the substrate binding pocket in influencing the substrate conformation and the resulting selectivity in BesD and HalB.

**Figure 6.**
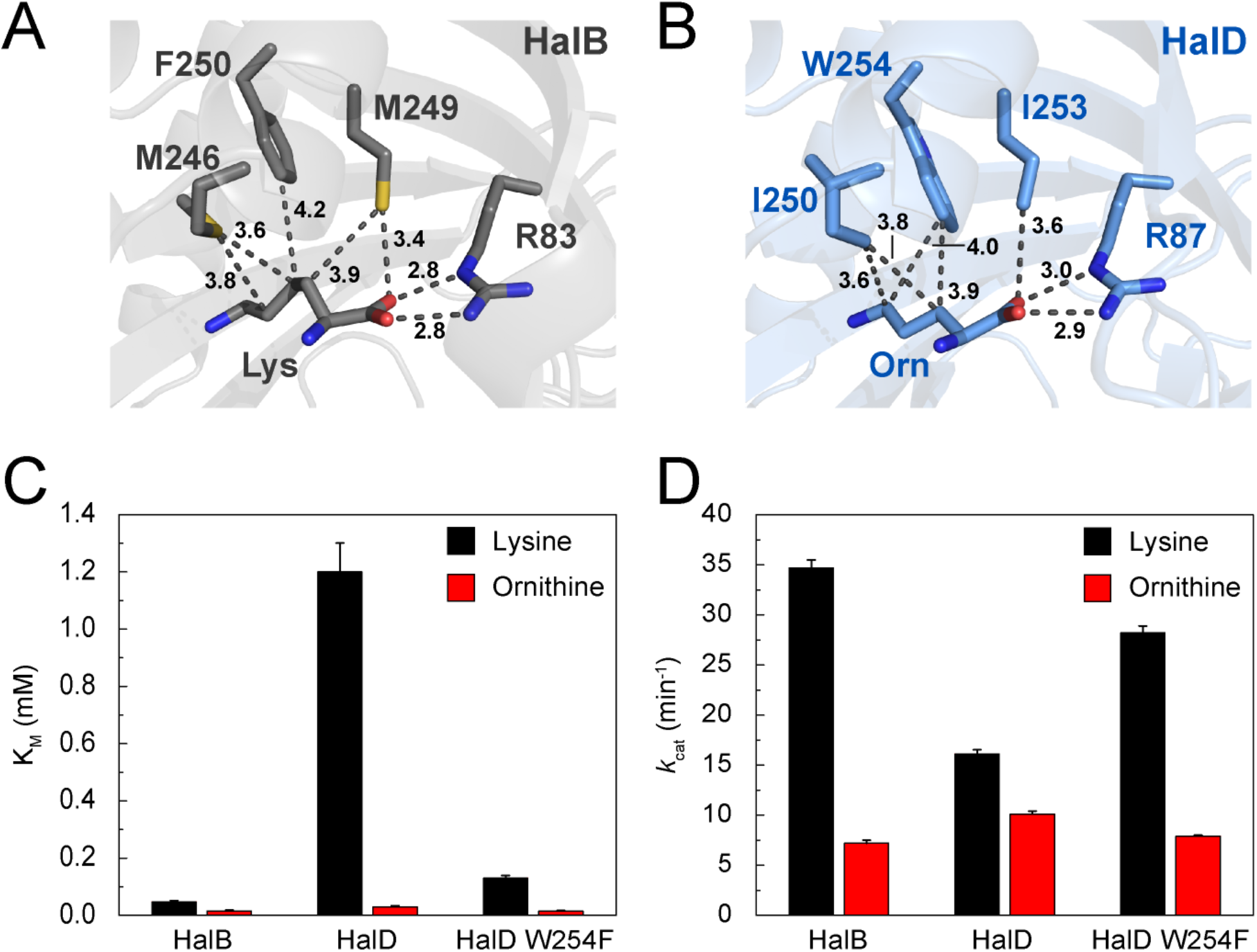
Engineering altered chain length selectivity. **(A)** In the lysine halogenase, HalB, the substrate binding pocket is composed of Met246, Met249, and Phe250. **(B)** In the ornithine halogenase, HalD, the substrate binding pocket is composed of Ile250, Ile253, and Trp254. **(C, D)** Steady-state kinetic analysis of HalB, HalD, and HalD W254F with lysine (black) or ornithine (red) as substrates. Michaelis-Menten data are in **Supplemental Figs. 15-17** as are mean ± s.d. (n = 3 technical replicates). The kinetic parameters *k*_*cat*_ and *K*_*M*_ were calculated by non-linear curve fitting to the Michaelis-Menten equation. Calculated *K*_*M*_ values are plotted as mean ± s.e. in panel (C). Calculated *k*_*cat*_ values are plotted as mean ± s.e. in (D).

We revisited the HalB crystal structure to determine whether additional residues may directly contribute to the observed regioselectivity. Phe250 in HalB is positioned 4.1 Å away from C_4_ and forms a pocket which stabilizes C_4_ away from the active site (**Fig. 3A**). To investigate the role of Phe250 on regioselectivity, we generated 4 separate NNK libraries at position 250 while keeping residue 246 constant as Met (wild type), Leu, Ile, or Val, the three mutations which yielded the greatest fluorescence in the screen. We screened over 150 colonies from each library using CuAAC (**Supplementary Figs. 8-11**). However, none of the mutations at Phe250 led to significantly increased yield of 4-chlorolysine beyond the original M246I, M246L, M246V point mutants (**Supplementary Tables 5-7**). These results suggest that point mutations are insufficient for fully altering regioselectivity, supporting the evolutionary logic of extensively remodelling the C terminal lid of BesD and HalB to achieve divergent outcomes.

### Crystal structure of ornithine halogenase HalD

In addition to members that make distinct regio-isomers of chlorolysine, the BesD amino acid halogenase family also contains members that display substrate selectivity. The enzyme HalD selectively chlorinates ornithine (k_cat_/K_M_ = 330 ± 70 mM^-1^min^-1^) over lysine (k_cat_/K_M_ = 13.2 ± 3.6 mM^-1^min^-1^)^60^. Interestingly, the two substrates only differ by one carbon. In HalD, the K_M_ for ornithine is 30 μM, which is within the range of intracellular ornithine concentrations^72^. In contrast, the lysine 4-halogenase HalA, a homolog that performs the same reaction as BesD, has a K_M_ for ornithine of 160 μM, which is well above the expected concentration of ornithine in the cell^72^. Although HalD chlorinates lysine and ornithine, the K_M_ value for lysine is 40 times higher (1.22 ± 0.12 mM vs 0.03 ± .004 mM), indicating a strong preference for ornithine^60^.

We initiated comparative structural studies to understand how HalD exhibits chain-length preference for the shorter ornithine substrate. We co-crystallized HalD with ornithine and solved the structure to 2.0 Å (**Supplementary Table 8**). The structure possessed clear electron density for the ornithine substrate within the active site (**Fig. 5A**). We also observed density for chloride, αKG, and a metal between the Fe-binding His149 and His 217 residues. Although iron is the catalytically-relevant metal, we assign the density in the structure as nickel based on our crystallography conditions and precedent for the persistence of nickel during nickel-affinity chromatography of Fe^II^/αKG-dependent enzymes^73^.

We next performed a structural comparison of HalD (ornithine 4-halogenase), BesD (lysine 4-halogenase), and HalB (lysine 5-halogenase). The cores of all three enzymes are structurally similar, and the substrates bind in a similar location in all three cases. Consistent with our proposed role of the C-terminal lid in dictating selectivity, the majority of the structural differences between the three enzymes occur within the C-termini (**Fig. 5B, Supplementary Fig. 12**). The C-terminal lid of the HalD aligns better with that of HalB (RMSD of 0.357 Å) than BesD (RMSD of 2.318 Å) (**Supplementary Fig. 13**). The lids of both HalD and HalB are alpha-helical in structure, while the C-terminus of BesD lacks secondary structure. Due to the high structural similarity of HalD and HalB, and because previous LC/MS results suggest that HalD produces 5-Cl-lysine rather than 4-Cl-lysine^60^, we chose HalB as the primary point of comparison for chain-length specificity.

A direct comparison of HalB and HalD revealed that the carboxylates and amines of lysine and ornithine are in very similar positions in the active site (shifted by 0.4, 0.3, and 0.5 Å for the α-carboxylate, α-amine, and ε-amine, **Fig. 5C**). However, while ornithine is bound in an extended conformation, the side chain of lysine is instead kinked away from the location of the catalytic Fe, making hydrophobic contacts with a binding pocket provided by the C-terminal covering lid. In the lysine halogenase, HalB, the substrate binding pocket is composed of Met246, Met249, and Phe250, which accommodate C_3_ and C_4_ of lysine (**Fig. 6A**). In contrast, the ornithine halogenase has a smaller pocket that is composed of Ile250, Ile253, and Trp254 (**Fig. 6B**). Notably, Trp254 in HalD is larger than Phe250 in HalB. As a result, the longer side chain of lysine is not readily accommodated in the HalD active site, consistent with the strong preference of HalD for ornithine.

### Engineering the HalD binding pocket to accommodate lysine

Although HalB and HalD are structurally similar, three key residues tailor the binding pocket to disfavor lysine binding in HalD. We therefore wondered whether expansion of the HalD binding pocket to resemble the pocket in HalB might enhance chlorination of lysine. As Trp254 appeared to have the most significant effect on the structure of the binding pocket, we cloned, expressed, and purified HalD W254F to expand the native substrate binding pocket. We next performed steady-state kinetic analyses of the wild type and mutant halogenase enzymes with lysine and ornithine as substrates (**Supplementary Figs. 14-16**). Consistent with our proposed role of the substrate binding pocket, mutagenesis of Trp254 to Phe lowered the K_M_ for lysine nearly 10-fold (0.13 mM) compared to wild type HalD (1.2 mM), approaching the K_M_ of wild type HalB (0.047 mM) (**Fig. 6C**). Mutagenesis also resulted in an increase in *k*_*cat*_ with lysine from 16 min^-1^ to 28 min^-1^, suggesting that expansion of the substrate binding pocket significantly improves accommodation of the lysine. Given that this single mutant resulted in nearly a complete switch of HalD parameters to match that of HalB, we next cloned, expressed, and purified HalD I250M I253M W254F to account for other differences in the active site in an attempt to further improve activity on lysine. While the triple mutant had slightly decreased K_M_ for lysine (0.45 mM), this mutant exhibited decreased *k*_*cat*_ for both ornithine and lysine suggesting that the additional mutations may destabilize the packing of the C-terminal lid onto the base of the structure, resulting in decreased rates (**Supplementary Fig. 17**). Taken together, these crystallographic and engineering studies suggest that mutagenesis of the C-terminal region enables engineering of amino acid halogenases with improved activity on target substrates.

## DISCUSSION

Biocatalytic C-H activation reactions takes advantage of the ability enzymes to achieve exquisite regio- and stereoselectivity in the generation of a substrate radical that can undergo subsequent functionalization with a nearby group. As such, engineering this reaction class can open the door to reimagining the interface between biological and chemical catalysis with the ultimate goal of achieving the synthesis of complex molecules at low cost and environmental impact. We have focused on studying the recently-discovered BesD family of amino acid halogenases, which directly chlorinate amino acids, as engineering the substrate scope of these enzymes could allow for metabolic and chemical diversification to produce an enormous range of structures.

Toward this goal, we have investigated the mechanisms by which enzymes within the BesD family evolved both regioselectivity and substrate selectivity with respect to site selection on a methylene chain. Halogenation has particularly stringent requirements for substrate positioning, explained by the strong distance dependence for chlorination compared to the thermodynamically-preferred competing hydroxylation pathway^43,44^. Unlike hydroxylation, rebound with chloride does not take place along the same reaction vector as C-H abstraction by Fe(IV)=O, so we therefore expect that this delicate balance of substrate positioning will be important in similar atom transfer reactions. In this work, we discovered that a variable substrate-binding lid that participates in an unprecedented conformational change featuring a helix-to-sheet transition upon substrate binding and is key to controlling reaction outcome. Interestingly, these lids are found in other Fe/αKG-dependent and Rieske enzymes and may play a similar role in substrate positioning in these enzymes given the shared need to close the active site and protect the substrate radical from water^62,74,75^.

Structural and biochemical comparisons of BesD (4-Cl-lysine), HalB (5-Cl-lysine) and HalD (4-Cl-ornithine) show that divergent regioselective outcomes arise by control of the substrate conformation within the active site rather than by translational shifting. A comparison of the lysine-bound structures of HalB and BesD reveals differences in the substrate covering lids that serve to orient the substrate to achieve the desired regioselectivity. In particular, Met246 shapes the substrate pocket of HalB such that C_5_ of lysine is most proximal to the catalytic site. Mutagenesis of Met246 to the smaller nonpolar residues Leu, Ile, and Val relieves the steric constraints and disrupts the regioselectivity, leading to halogenation at C_4_. Consistent with this model, the substrate covering lid of HalD enables selectivity for the smaller, 5-carbon substrate, ornithine, over the larger 6-carbon substrate, lysine. While the covering lid of HalD is structurally similar to that of HalB, the identity of residues that form the substrate binding pocket vary. Specifically, HalD contains the larger Trp254 compared to Phe250 found in HalB and fails to accommodate lysine as effectively as it does ornithine (40-fold difference in K_M_). The structure-guided expansion of the HalD active site to accommodate lysine with a K_M_ rivaling that of native HalB supports the role of the C-terminal lid in substrate selection and sets the stage for future engineering efforts to harness the reactivity of the iron complex for C–H activation and halogenation of non-native substrates.

The enzymes in this study showcase the tremendous power of biocatalysis. This work provides fundamental insights into how BesD, HalB, and HalD use the same catalytic metal center to perform subtly different reactions with high selectivity. In addition to chlorination, enzymes in this family have been shown to perform bromination and azidation, further illustrating the versatility of these enzymes for biocatalysis.^60^ The application and evolution of radical halogenases toward modification of C-H bonds has the potential to bridge biological and chemical methods for bond formation, broadening the scope of accessible compounds.

## Acknowledgments

This work was funded by generous support from the National Institutes of Health to M.C.Y.C. (R01 GM134271). E.N.K. acknowledges the support of a National Institutes of Health NRSA Training Grant (1 T32 GMO66698). M.E.N. acknowledges the support of a National Science Foundation Graduate Research Fellowship. C.V.S. acknowledges the support of a Berkeley Fellowship for Graduate Study. J.A.M. acknowledges the support of a UC Berkeley Chancellor’s Fellowship, Howard Hughes Medical Institute Gilliam Fellowship, and National Institutes of Health NRSA Training Grant (1 T32 GMO66698). X-ray data were collected at the Advanced Light Source Beamline 8.3.1, which is operated by the University of California Office of the President, Multicampus Research Programs and Initiatives (MR-15-328599), the National Institutes of Health (R01 GM124149 and P30 GM124169), Plexxikon Inc., and the Integrated Diffraction Analysis Technologies program of the U.S. Department of Energy Office of Biological and Environmental Research. The Advanced Light Source is a national user facility operated by Lawrence Berkeley National Laboratory on behalf of the U.S. Department of Energy under contract number DEAC02-05CH11231, Office of Basic Energy Sciences. The funds for the 900 MHz NMR spectrometer housed in the QB3 Institute in Stanley Hall at University of California, Berkeley were kindly provided by the NIH (GM68933).

## Author contributions

E.N.K and M.E.N performed enzyme characterization experiments, library generation, screen development, and library analysis, and protein crystallography. K.H.S. and C.V.S. performed protein crystallography. N.A.S. assisted with enzyme characterization and screening experiments. J.A.M. and D.C.M. contributed to screen development. E.N.K., M.E.N., and M.C.Y.C. wrote the paper with contributions from all other authors.

