## Supplementary Information for "Regioselective control of biocatalytic C–H activation and halogenation"

#### Supplementary Methods

*Commercial materials* S3
*Sequence alignment of BesD, HalB, and HalD* S3
*Bacterial strains* S4
*Construction of plasmids* S4
*Screening the HalB library for alkyne formation* S4
*Expression of $His_{10}$-tagged proteins* S5
*Purification of $His_{10}$-tagged proteins* S5
*Preparation of proteins for crystallization* S6
*HalB crystallization and data collection* S6
*HalD crystallization and data collection* S6
*Structure determination* S6
*Steady-state kinetic analysis of HalB variants* S7
*Procedure for LC/MS analysis of polar metabolites* S7
*In vitro LC/MS assay of halogenase variants* S7

#### Supplementary Results

**Supplementary Table 1.** *Strains, plasmids, oligonucleotides, and synthetic gene sequences* S9
**Supplementary Figure 1.** *Sequence alignment of BesD, HalA, HalB, and HalD* S12
**Supplementary Table 2.** *Crystallography table of HalB with glycine bound* S13
**Supplementary Table 3.** *Crystallography table of HalB with lysine bound* S14
**Supplementary Figure 2.** *Mechanism of $Fe^{II}/\alpha$KG-dependent halogenases* S15
**Supplementary Figure 3.** *CuAAC reaction scheme* S16
**Supplementary Figure 4.** *M246NNK CuAAC screening results* S17
**Supplementary Table 4.** *M246NNK sequencing results* S18
**Supplementary Figure 5.** *Quantification of propargylgycine production by CuAAC* S19
**Supplementary Figure 6.** *Kinetics of purified M246 variants* S20
**Supplementary Figure 7.** *LCMS quantification of 5-Cl-lysine production by HalB variants* S21
**Supplementary Figure 8.** *F250NNK CuAAC screening results* S22
**Supplementary Figure 9.** *M246L F250NNK CuAAC screening results* S23
**Supplementary Table 5.** *M246L F250NNK sequencing results* S24
**Supplementary Figure 10.** *M246V F250NNK CuAAC screening results* S25
**Supplementary Table 6.** *M246V F250NNK sequencing results* S26
**Supplementary Figure 11.** *M246I F250NNK CuAAC screening results* S27

**Supplementary Table 7.** *M246I F250NNK sequencing results* S28
**Supplementary Table 8.** *Crystallography table of HalD* S29
**Supplementary Figure 12.** *Predicted stereochemistry of hydrogen atom abstraction* S30
**Supplementary Figure 13.** *Overall fold comparison of HalB, BesD, and HalD* S31
**Supplementary Figure 14.** *Active site comparison of HalD and BesD* S32
**Supplementary Figure 15.** *SDS-PAGE gel of purified proteins used in the study* S33
**Supplementary Figure 16.** *Kinetics of HalB on lysine and ornithine* S34
**Supplementary Figure 17.** *Kinetics of HalD W254F* S35
**Supplementary Figure 18.** *Kinetics of HalD I250M I253M W254F* S36

**References** S37

#### Materials and Methods

**Commercial materials.** Luria-Bertani (LB) Broth Miller, LB Agar Miller, Terrific Broth (TB), and glycerol were purchased from EMD Biosciences (Darmstadt, Germany). Carbenicillin (Cb), kanamycin (Km), chloramphenicol (Cm), isopropyl-β-D-thiogalactopyranoside (IPTG), sodium chloride, dithiothreitol (DTT), 4-(2-hydroxyethyl)-1-piperazineethanesulfonic acid (HEPES), magnesium chloride hexahydrate, acetonitrile, ethylene diamine tetraacetic acid disodium dihydrate (EDTA), hydrochloric acid, 3kDa MWCO dialysis tubing, ammonium bicarbonate, sodium acetate, chloroform, and sodium hydroxide were purchased from Fisher Scientific (Pittsburgh, PA). Phosphoenolpyruvate (PEP), adenosine triphosphate sodium salt (ATP), nicotinamide adenine dinucleotide reduced form dipotassium salt (NADH), pyruvate kinase, lactate dehydrogenase, lysozyme, poly(ethyleneimine) solution (PEI), ammonium iron (II) sulfate hexahydrate, α-ketoglutaric acid sodium salt, β-mercaptoethanol (βME), N,N,N',N'-tetramethyl-ethane-1,2-diamine (TEMED), betaine, dimethylsulfoxide (DMSO), sodium L-ascorbate, acetonitrile (LC/MS-grade), ammonium formate (LC/MS-grade), Deuterium oxide, L-lysine for crystallography, 2-(N-Morpholino)ethanesulfonic acid hydrate, 4-morpholineethanesulfonic acid (MES), PEG 3350, PEG 1500, PEG 550 MME, Pall Nanosep centrifugal Omega membrane (3,000 MWCO), AeraSeal films, and copper sulfate ($CuSO_4$) were purchased from Sigma-Aldrich (St. Louis, MO). PageRuler Plus Prestained Protein Ladder was purchased from ThermoFisherScientific (Waltham, MA). Succinyl-CoA synthetase was purchased from Megazyme International (Bray, Ireland). Lysine 2HCl ($^{13}C_6$, 99%; $^{15}N_2$, 99%), and 4,4,5,5,-$D_4$-lysine were purchased from Cambridge Isotope Laboratories (Andover, MA). Formic acid was purchased from Acros Organics (Morris Plains, NJ). Restriction enzymes, T4 DNA ligase, Phusion DNA polymerase, T5 exonuclease, and Taq DNA ligase, were purchased from New England Biolabs (Ipswich, MA). Deoxynucleotides (dNTPs), were purchased from Invitrogen (Carlsbad, CA). Oligonucleotides were purchased from Integrated DNA Technologies (Coralville, IA), resuspended at a stock concentration of 100 µM in water and stored at either 4 °C for immediate usage or -20 °C for long-term storage. DNA purification kits and Ni-NTA agarose were purchased from Qiagen (Valencia, CA). Zirconia/silica beads were purchased from BioSpec Products (Bartlesville, OK). Complete EDTA-free protease inhibitor was purchased from Roche Applied Science (Penzberg, Germany). PD-10 desalting columns and the Superdex 75 16/60 pg column were purchased from GE Healthcare (Pittsburg, PA). Amicon Ultra 10,000 MWCO centrifugal concentrators and Milli-Q Gradient water purification system were purchased from Millipore (Billerica, MA). Acrylamide/bis-acrylamide (30%, 37.5:1), electrophoresis grade sodium dodecyl sulfate (SDS), and ammonium persulfate were purchased from Bio-Rad Laboratories (Hercules, CA). Ultrayield baffled flasks were purchased from Thompson Instrument Company (Oceanside, CA). Tris-hydroxypropyltriazolylmethylamine (THPTA) was purchased from Click Chemistry Tools (Scottsdale, AZ). 96-well plates with 2 ml volume and v-bottom were purchased from Corning (Corning, NY).

**Sequence alignment of BesD, HalB, and HalD.** Sequences of SlBesD (WP_030791981), ScBesD (WP_014151497), SwHalB (SDN46247), PkHalD (WP_046063366), and PfHalA (WP_016975823) were aligned using MUSCLE [*1*], which was also used to compute the percent identity matrices.

**Bacterial strains.** *E. coli* DH10B-T1$^R$ was used for plasmid construction. BL21(DE3)–Star harboring the pRARE2 plasmid was used for heterologous protein production of all halogenase variants.

**Construction of plasmids.** Gibson assembly [*2*] was used to carry out plasmid construction using *E. coli* DH10B-T1$^R$ as the cloning host. PCR amplifications were carried out with Phusion polymerase or Q5 polymerase (New England Biolabs, Ipswich MA) using the oligonucleotides listed in **Table 1B**. GeneBlock sequences are listed in **Table 1C** along with primers used to construct plasmids. Following plasmid construction, all cloned inserts were sequenced at Azenta Life Sciences (San Francisco, CA).

*Plasmids for protein expression.* The intermediate cloning plasmid, pET16-His-PrescissionCutSite-IMPDH, was constructed by amplification of IMPDH from *E. coli* gDNA followed by insertion into NdeI/BamHI-digested pET16b. The halogenase WP_014151497 (*S. cattleya* BesD) was amplified from genomic DNA of the corresponding organism and inserted into NdeI/BamHI-digested pET16-$His_{10}$-PrescissionCutSite-IMPDH. WP_046063366 (*P. kilonensis* HalD) was obtained as a GeneBlock and amplified with primers as noted (**Table 1B**) before Gibson insertion into NdeI/BamHI-digested $His_{10}$-PrescissionCutSite-IMPDH. WP_030791981 (*S. lavanduligriseus* BesD) and SDN46247 (*S. wuyuanensis HalB*) were obtained as GeneBlocks containing the overhangs required for direct Gibson insertion into NdeI/BamHI-digested pET16-His-Prescission-IMPDH.

*Plasmids for library generation and screening.* To clone the M246NNK library of SwHalB, SwHalB was amplified in two pieces from the pET16-His-SwHalB plasmid. The first piece was amplified with SwHalB-F and SwHalB246NNK-R. The second piece was amplified with SwHalB246NNK-F and pET16-anneal-R. Then both pieces were inserted into NdeI/HindIII-digested pET16-$His_{10}$-PrescissionCutSite-IMPDH to yield the mutant constructs. The F250NNK, M246V F250NNK, M246L F250NNK, and M246I F250NNK libraries were cloned using SwHalB, SwHalB M246V, SwHalB M246L, and SwHalB M246I, respectively, as templates. The first piece was amplified with SwHalB-F and the corresponding reverse primers. The second piece was amplified with pET16-anneal-R and the corresponding forward primers. Both pieces were inserted into NdeI/HindIII-digested pET16-$His_{10}$-PrescissionCutSite-IMPDH to yield the mutant constructs. To generate pBesCB, a fragment containing besC-T7p-MBP-besB was PCR amplified from pPra [*3*] using primers pBesCB-F/R and cloned into pACYC-DUET using Gibson ligation and NcoI/KpnI cut sites. Inserts for cloning the chimeric fusion of BesD and HalB (ChimeraDB) were amplified in four pieces from pET16-His-SlBesD (pieces 1-3) or pET16-His-SwHalB (piece 4) using the ChDB primers in **Table 1B**. Pieces 1-3 were annealed using overlap extension PCR [*4*]. The fused PCR product, along with piece 4, was inserted into NdeI/HindIII-digested pET16-$His_{10}$-PrescissionCutSite-IMPDH to yield the chimeric construct.

**Screening the HalB library for alkyne formation.** Chemically-competent *E. coli* BL21 cells harboring the pBesCB plasmid ($Cm^R$) were transformed with the SwHalB NNK library plasmids ($Cb^R$) and grown on LB plates with the appropriate antibiotic. Colonies were inoculated into individual wells of a 96-well plate containing 200 μL LB with the appropriate antibiotic and grown overnight. Dense cultures were diluted 20-fold into 96-well plates (2 ml v-bottom, Corning, NY) containing 1 ml of LB, covered with Aeraseal Film (Sigma) and grown to mid-log phase (OD = 0.6 to 0.8) before chilling the plates on ice for 30 min. Cells were pelleted by centrifugation at 4,000 × *g* for 20 min and then resuspended in 1 ml of M9 media (sodium phosphate (6.6 g/L), potassium phosphate (3 g/L), sodium chloride (0.5 g/L), ammonium chloride (1 g/L), magnesium sulfate (2 mM), calcium chloride (0.1 mM), glucose (4% (*w/v*)), thiamine (1 mg/L)) supplemented with 0.5 mM lysine, 0.25 mM IPTG, and the appropriate antibiotics. Cells were grown for 2 d at 16°C with shaking at 225 rpm. After 2 d of growth in M9 media, the cultures were supplemented with lysine (0.2 mM), sodium αKG (0.2 mM), and NaCl (0.2 mM) and grown for 2 additional days. Note that at the end of the day 4, some of the cells had settled in the bottom of the plate, despite shaking during growth. The remaining cells were then pelleted by centrifugation for 15 min at 4,000 × *g*. Media samples (50 µL) containing Pra were pipetted into a 96 well black plate with a clear bottom (Corning, NY). To initiate the reaction, 50 µL of a 2× CuAAC reaction solution in 1× PBS was added to each well to give the final reaction solution (CalFluor 488 azide (1 µM), copper (II) sulfate (0.5 mM), BTTAA (0.1 mM), and sodium ascorbate (5 mM)). Reactions were performed for 30 min in the dark before analyzing fluorescence of the Pra-CalFluor product ($\lambda_{ex}$ = 485 nm, $\lambda_{em}$ = 528 nm) using a SynergyMx Microplate Reader (BioTek) at room temperature. Cells from the fluorescent wells were grown overnight, miniprepped (Qiagen), and sequenced with the Prescission-seq-F primer in **Table 1B**, which anneals to the SwHalB NNK vector but not to the pBesCB vector which was also present in the cells. To quantify the production of Pra by the wild type BesD, HalB, and M246 mutants, we produced a standard curve of Pra click fluorescence in the spent media. Samples of a lysine hydroxylase were grown according to the protocol above. On day 4, 12 10x stock solutions of Pra ranging from 2 µM to 180 µM were prepared. After removal of the cells by centrifugation, 40 µl of spent media was added to 10 µl of 10x Pra solution before the sample was added to 50 µl of the 2x CuAAC reaction solution. After collection of fluorescence data, a linear range of Pra concentrations was selected from 0.2 to 7 µM.

**Expression of His$_{10}$-tagged proteins.** *E. coli* BL21 Star (DE3) harboring the pRARE2 plasmid was transformed with the appropriate protein expression plasmid. An overnight TB culture of the freshly transformed cells was used to inoculate TB (1 L) containing the appropriate antibiotics (50 µg/ml carbenicillin or kanamycin with 50 µg/ml chloramphenicol) in a 2.8 L-baffled shake flask to $OD_{600}$ = 0.05. The cultures were grown at 37°C at 200 rpm to $OD_{600}$ = 0.6 to 0.8 at which point cultures were cooled on ice for 20 min, followed by induction of protein expression with IPTG (0.25 mM) and overnight growth at 16°C. Cell pellets were harvested by centrifugation at 9,800 × *g* for 7 min at 4ºC and stored at -80ºC.

**Purification of His$_{10}$-tagged proteins.** Frozen cell pellets were thawed and resuspended at 5 ml/g of cell paste in lysis buffer (50 mM HEPES, 300 mM NaCl, 10 mM imidazole, 20 mM βME, 20% (*v/v*) glycerol, pH 7.5) supplemented with EDTA-free Protease Inhibitor Cocktail (Roche). The cell paste was homogenized and then lysed by sonication with a Qsonica Q700 sonicator (Amplitude = 45, 5 s on, 25 s off, 2.5 min total process time, ½" tip). The lysate was then centrifuged at 13,500 × *g* for 20 min at 4°C to separate the soluble and insoluble fractions. DNA was precipitated in the soluble fraction with 0.15% (*w/v*) polyethyleneimine and stirring at 4°C for 30 min. The precipitated DNA was then removed by centrifugation at 13,500 × *g* for 20 min at 4°C. The soluble lysate was incubated with Ni-NTA (0.5 ml resin/g of cell paste) for 45 min at 4°C, then resuspended and loaded onto a column by gravity flow. The column was washed with wash buffer (50 mM HEPES, 300 mM NaCl, 20 mM imidazole, 20 mM βME, 20% (*v/v*) glycerol, pH 7.5) for 15-20 column vol. The column was then eluted with elution buffer (50 mM HEPES, 300 mM NaCl, 300 mM imidazole, 20 mM βME, 20% (*v/v*) glycerol, pH 7.5). Fractions containing the target protein were pooled by $A_{280\ nm}$ and concentrated using an Amicon Ultra spin concentrator (10 kDa MWCO, Millipore). Protein was then exchanged into storage buffer (50 mM HEPES, 100 mM sodium chloride, 20% (*v/v*) glycerol, 1 mM DTT, pH 7.5) using PD-10 desalting columns. Final protein concentrations before storage were estimated using the $\varepsilon_{280\ nm}$ calculated by ExPASy ProtParam [5] and measured by nanodrop. They are as follows: SwHalB: 10.67 mg/ml ($\varepsilon_{280\ nm}$ = 45,045 $M^{-1}\ cm^{-1}$), SwHalB M246L: 3.93 mg/ml ($\varepsilon_{280\ nm}$ = 45,045 $M^{-1}\ cm^{-1}$), SwHalB M246I: 4.35

mg/ml ($\varepsilon_{280\ nm}$ = 45,045 $M^{-1}$ $cm^{-1}$), SwHalB M246V: 4.35 mg/ml ($\varepsilon_{280\ nm}$ = 45,045 $M^{-1}$ $cm^{-1}$), SlBesD: 6.48 mg/ml ($\varepsilon_{280\ nm}$ = 50,545 $M^{-1}cm^{-1}$), SwHalB F250W: 5.73 mg/ml ($\varepsilon_{280\ nm}$ = 50,420 $M^{-1}$ $cm^{-1}$), SwHalB M246I M249I F250W: 5.97 mg/ml ($\varepsilon_{280\ nm}$ = 50,420 $M^{-1}$ $cm^{-1}$), PkHalD: 15 mg/ml ($\varepsilon_{280\ nm}$ = 60,390 $M^{-1}$ $cm^{-1}$), PkHalD W254F: 5.74 mg/ml ($\varepsilon_{280\ nm}$ = 54,890 $M^{-1}$ $cm^{-1}$), PkHalD I250M I253M W254F: 4.14 mg/ml ($\varepsilon_{280\ nm}$ = 54,890 $M^{-1}$ $cm^{-1}$). All proteins were aliquoted, flash-frozen in liquid nitrogen, and stored at -80°C.

**Preparation of proteins for crystallization.** Following elution from the Ni-NTA column, the proteins were incubated with Prescission protease (1 mg protease/50 mg protein) and dialyzed 1:50 overnight into dialysis buffer to remove imidazole. Cleaved and dialyzed protein was passed through Ni-NTA (2 ml) to remove Prescission and the $His_{10}$ tag. The eluent was the diluted to a final salt concentration of 20 mM NaCl using Buffer A (50 mM HEPES, 20% (*v/v*) glycerol, 1 mM DTT, 1 mM EDTA, pH 7.5) and loaded onto a 5 ml HiTrap-Q column for ion exchange with the AKTA Purifier FPLC system (GE Healthcare). The protein was eluted using a gradient from 0-100% buffer A to buffer B (50 mM HEPES, 1 M NaCl, 20% (*v/v*) glycerol, 1 mM DTT, 1 mM EDTA, pH 7.5) over 40 min. In the case of PkHalD, EDTA was omitted from the ion exchange purification procedure. The protein sample was concentrated to 2 ml and loaded onto a HiLoad Superdex 75 16/600 pg (GE Healthcare) column equilibrated with SEC Buffer (20 mM HEPES, 100 mM NaCl, 1 mM DTT, pH 7.5). The protein eluent was concentrated to 15 mg/ml and glycerol was added to a final concentration of 5% (*v/v*) before flash freezing in liquid nitrogen and storage at -80°C.

**HalB crystallization and data collection (glycine bound).** SwHalB crystals were obtained by the hanging drop vapor diffusion method by combining equal volumes of protein solution (SwHalB (2.5 mg/ml), lysine (3 mM), αKG (3 mM, pH 7)) and reservoir solution (succinate phosphate glycine (SPG) buffer (100 mM), 18% (w/v) PEG 1500, pH 5.2). The 1 M SPG buffer stock contained succinic acid (125.3 mM), sodium dihydrogen phosphate monohydrate (437.7 mM), and glycine (436.9 mM). Crystals grew in three days and were flash frozen in liquid nitrogen. Data were collected at Beamline 8.3.1 at the Advanced Light Source (Lawrence Berkeley National Laboratory) at a wavelength of 1.11 Å.

**HalB crystallization and data collection (lysine bound).** Crystals of HalB from *S. wuyuanensis* (SwHalB) were obtained by the hanging drop vapor diffusion method by combining equal volumes of protein solution (SwHalB (3 mg/ml), lysine (3 mM), αKG (3 mM, pH 7)) and reservoir solution (succinate phosphate lysine (SPL) buffer (100 mM), 20% (*w/v*) PEG 1500, pH 5.5). The 1 M SPL buffer stock was made as a modification of SPG (succinate phosphate glycine) buffer and contained succinic acid (125.3 mM), sodium dihydrogen phosphate monohydrate (437.7 mM), and lysine (436.9 mM). Crystals grew in three days and were transferred to an Eppendorf tube containing 250 μL of reservoir solution and vortexed for 30 seconds with 10 × 1 mM diameter zirconia/silica beads to produce a micro-seed solution. The seed solution was stored at -80°C for future use. Subsequent crystals were prepared by micro-seeding equal volumes of protein solution (2 mg/ml) SwHalB, lysine (3 mM), αKG (3 mM, pH 7)) and reservoir solution (succinate phosphate lysine (SPL) buffer, pH 5.4 (100 mM), 20% (*w/v*) PEG 1500). Crystals grew in one day and were flash frozen in liquid nitrogen. Data were collected at Beamline 8.3.1 at the Advanced Light Source (Lawrence Berkeley National Laboratory) at a wavelength of 1.11 Å.

**HalD crystallization and data collection.** Crystals of HalD from *P. kilonensis* (HalD) were obtained by the hanging drop vapor diffusion method by combining equal volumes of protein solution (HalD (6 mg/ml), ornithine (3 mM), αKG (3 mM, pH 7)) and reservoir solution (MES buffer (100 mM), ~35% (*w/v*) PEG 550 MME, pH 6.3). Crystals grew within two days and microseeds were generated as described above for HalB. Subsequent crystals were obtained by microseeding equal volumes of protein solution (HalD (6 mg/ml), ornithine (3 mM), αKG (3 mM, pH 7)) and reservoir solution (MES buffer (100 mM), 27% (*w/v*) PEG 550 MME, pH 6.2). Crystals grew in one day and were flash frozen in liquid nitrogen. Data were collected at Beamline 8.3.1 at the Advanced Light Source (Lawrence Berkeley National Laboratory) at a wavelength of 1.11 Å.

**Structure determination.** Data were processed with XDS [*6*] and scaled and merged with Aimless [*7*]within the CCP4 suite [*8*]. The data were phased with molecular replacement using the structure of BesD (6NIE) as a search model. The structures were refined iteratively in COOT [*9*] and Phenix [*10*]. Ligands were added to the model using COOT and refined in Phenix. Omit maps for the ligand complex including were generated using the Phenix Map function by removing atoms of interest from refinement prior to calculation of the maps. The structure was analyzed in Pymol, which was also used to create figures.

**Kinetic analysis of HalB variants.** Steady-state kinetic analysis was performed by monitoring NADH consumption through a coupled assay [*11*]. Reactions (100 µL) contained ATP (2.5 mM), $MgCl_2$ (5 mM) phosphoenol-pyruvate (PEP, 1 mM), NADH (0.3 mM) lactate dehydrogenase (LDH, 10 U/ml), pyruvate kinase (PK, 10 U/ml) succinyl-CoA synthetase (SCS, 3.2 U/ml), coenzyme A (1 mM), sodium αKG (1 mM), $(NH_4)_2Fe(SO_4)_2 \cdot 6H_2O$ (0.2 mM), sodium chloride (10 mM), and sodium ascorbate (2 mM) in 100 mM HEPES buffer (pH 7.5). Reactions were initiated by addition of the halogenase variant (2.5 µM) in the presence of varying concentrations of L-lysine · HCl (0 - 1 mM). Initial rates of NADH consumption were measured by monitoring $A_{340}$ using a SpectraMax M2 Microplate Reader (Molecular Devices) at room temperature. $k_{cat}$ and $K_M$ were determined by fitting to initial rate data with Origin (OriginLab, Northampton, MA) using the equation:

$$v_0 = \frac{k_{cat}[S]}{K_M + [S]}$$

where $v_o$ is the initial rate and [*S*] is the substrate concentration. For data in which substrate inhibition was observed, $k_{cat}$, $K_M$, and $K_i$ were determined using the following equation.

$$v_0 = \frac{k_{cat}[S]}{K_M + [S](1 + [S]/Ki)}$$

**General procedure for high resolution HPLC/MS analysis of polar metabolites with HILIC.** Samples containing polar metabolites were analyzed using an Agilent 1290 UPLC-QTOF on a SeQuant ZIC-pHILIC (5 µm, 2.1 × 100 mm; EMD-Millipore) using the following buffers: Buffer A (90% acetonitrile, 10% water, 10 mM ammonium formate) and Buffer B (90% water, 10% acetonitrile, 10 mM ammonium formate). A linear gradient from 95% to 60% Buffer A over 17 min followed by a linear gradient from 60% to 33% Buffer A over 8 min was then applied at a flow rate of 0.2 ml/min. Mass spectra were acquired in positive ionization scan mode using a 6530 QTOF (Agilent).

***In vitro* assays of halogenase variants.** Reactions (50 µL) contained L-lysine · HCl (3 mM), sodium αKG (5 mM), sodium ascorbate (5 mM), $(NH_4)_2Fe(SO_4)_2 \cdot 6H_2O$ (1 mM), and sodium chloride (5 mM) in 100 mM HEPES buffer (pH 7.5). Reactions were initiated by addition of purified SwHalB variants (20 µM final concentration) and allowed to proceed for 25 min at room temperature before quenching in 2 vol of methanol with 1% (*v/v*) formic acid. Following centrifugation at 18,000 × *g* for 20 min (4 °C) to remove precipitated protein, samples were analyzed by LC/MS on an Agilent 1290 UPLC-QTOF using the protocol for polar metabolite analysis above.

### Supplementary Results

**Supplementary Table 1. Strains, plasmids, oligonucleotides, and synthetic gene sequences.** (A) Strains and plasmids used in this study. (B) Oligonucleotides used for plasmid construction and sequencing. (C) Sequences of synthetic genes.

#### A. Strains and plasmids

| Strain | Description | Source |
|---|---|---|
| BL21 Star (DE3) | *F-ompT hsdSB (rB-, mB-) gal dcm rne131* (DE3) | Thermo-Fisher |

| Plasmid | Description | Source |
|---|---|---|
| pET16b-His$_{10}$-SlavenduligriseusBesD | His$_{10}$-SlavenduligriseusBesD (T7), *lacI*, Cb$^R$, ColE1 | This study |
| pET16b-His$_{10}$-SwuyuanensisHalB | His$_{10}$-SwuyuanensisHalB (T7), *lacI*, Cb$^R$, ColE1 | This study |
| pET16b-His$_{10}$-SwHalB M246NNK | His$_{10}$-SwHalB M246NNK (T7), *lacI*, Cb$^R$, ColE1 | This study |
| pET16b-His$_{10}$-SwHalB F250NNK | His$_{10}$-SwHalB M246NNK (T7), *lacI*, Cb$^R$, ColE1 | This study |
| pET16b-His$_{10}$-SwHalB M246L F250NNK | His$_{10}$-SwHalB M246L F250NNK (T7), *lacI*, Cb$^R$, ColE1 | This study |
| pET16b-His$_{10}$-PrescissionCutSite-IMDPH | His$_{10}$-PrescissionCutSite-IMDPH (T7), lacI, Cb$^R$, ColE1 | This study |
| pET16b-His$_{10}$-PkHalD W254F | His$_{10}$-PkilonensisHalD (T7), *lacI*, Cb$^R$, ColE1 | This study |
| pET16b-His$_{10}$-PkHalD I250M I253M | His$_{10}$-PkHalD W254F (T7), *lacI*, Cb$^R$, ColE1 | |
| W254F | His$_{10}$-PkHalD W254F (T7), *lacI*, Cb$^R$, ColE1 | |
| pET16b-His$_{10}$-SwHalB M246V F250 NNK | His$_{10}$-PkHalD I250M I253M W254F (T7), *lacI*, Cb$^R$, ColE1 | |
| pET16b-His$_{10}$-SwHalB M246I F250 NNK | His$_{10}$-SwHalB M246V F250NNK (T7), *lacI*, Cb$^R$, ColE1 | |
| pET16b-His$_{10}$-SwHalB M246I M249I | His$_{10}$-SwHalB M246I F250NNK (T7), *lacI*, Cb$^R$, ColE1 | |
| F250W | His$_{10}$-SwHalB M246I M249I F250W (T7), *lacI*, Cb$^R$, | |
| pET16b-His$_{10}$-SwHalB F250W | ColE1 | |
| | His$_{10}$-SwHalB F250W (T7), *lacI*, Cb$^R$, ColE1 | |
| pACYC-DUET-BesC.MBP-BesB | BesC (T7) and MBP-BesB (T7) from *P. fluorescens*, *lacI*, Cm$^R$, ColE1 | This study |
| pPra | BesD (T7) and BesC (T7) from *P. fluorescens*, MBP-sBesB (T7) from *S.* sp. NRRL S-1448, *lacI*, Cb$^R$, ColE1 | Ref. 3 |
| pACYC-DUET | *lacI*, f1, Cm$^R$, ColE1 | Novagen |
| pRARE2 | tRNA$^{Ile}$(AUA), tRNA$^{Ser}$(AGG), tRNA$^{Ser}$(AGA), tRNA$^{Asp}$(CUA), tRNA$^{Gly}$(CCC), tRNA$^{Gly}$(GGA), Cm$^R$ | Novagen |

#### B. Oligonucleotide sequences

| Name | Sequence |
|---|---|
| His$_{10}$-SlBesD-F | gcggccatctagaagtgctttttcagggcccgcatatgagcagcaatgggcaggaaagcaccgtcgtca |
| His$_{10}$-SlBesD-R | cctttcgggctttgttagcagccggatcctcgagtcagtcatccagtgcagcggctgccggc |
| His$_{10}$-SwHalB-F | agcagcggccatctagaagtgctttttcagggcccgcatatgaccgagaacagcatcaccgc |
| His$_{10}$-SwHalB M246NNK-R | cagcttcctttcgggctttgttagcagccggatcctcatcagtcgaacatcgcgttmnnggtctcgtgggtggtcgcc |
| His$_{10}$-SwHalB F250NNK-R | cagcttcctttcgggctttgttagcagccggatcctcatcagtcmnncatcgcgttcatggtctcgtgggtggtcgcc |
| His$_{10}$-SwHalB M246V F250NNK-R | cagcttcctttcgggctttgttagcagccggatcctcatcagtcmnncatcgcgttaacggtctcgtgggtggtcgcc |
| His$_{10}$-SwHalB M246L F250NNK-R | cagcttcctttcgggctttgttagcagccggatcctcatcagtcmnncatcgcgttcaaggtctcgtgggtggtcgcc |
| His$_{10}$-SwHalB M246I F250NNK-R | cagcttcctttcgggctttgttagcagccggatcctcatcagtcmnncatcgcgttaatggtctcgtgggtggtcgcc |
| His$_{10}$-SwHalB F250W-F | agaccatgaacgcgatgtgggactgatgaggatccggct |
| His$_{10}$-SwHalB F250W-R | agccggatcctcatcagtcccacatcgcgttcatggtct |

| | |
|---|---|
| $His_{10}$-SwHalB M246I M249I F250W-F | gcgaccacccacgagaccatcaacgcgatctgggactgatgaggatccggctgct |
| $His_{10}$-SwHalB M246I M249I F250W-R | agcagccggatcctcatcagtcccagatcgcgttgatggtctcgtgggtggtcgc |
| pET16-anneal-R | cgtatcacgaggccctttcgtcttc |
| $His_{10}$-PkilonensisHalD-F | agcagcggccatctagaagtgctttttcagggcccgcatatgagcgaacaaacgtccagcttg |
| $His_{10}$-PkilonensisHalD-R | ttcctttcgggctttgttagcagccggatcctcgagtcaggcttcttcggtggtccg |
| $His_{10}$-PkHalD W254F-F | aaagcatcgaagtgatcttcgacaccaaggcccggacca |
| $His_{10}$-PkHalD W254F-R | tggtccgggccttggtgtcgaagatcacttcgatgcttt |
| $His_{10}$-PkHalD I250M I253M W254F-F | cgaaagcatggaagtgatgttcgacaccaaggcccggacca |
| $His_{10}$-PkHalD I250M I253M W254F-R | ttggtgtcgaacatcacttccatgctttcgtgggcgacatcgg |
| ChDB1-F | gcggccatctagaagtgctttttcagggcccgcatatgagcagcaatgggcaggaaagcaccgtcgtca |
| ChDB1-R | ctggaggcggagatcaatccgctcggcgtatttctccagcagccggtc |
| ChDB2-F | aatacgccgagcggattgatctccgcctccagaccaccggatacacccg |
| ChDB2-R | gataatgaattcctcgtcggcccaggggcaggggtgcagtttctcgccgg |
| ChDB3-F | cccctgggccgacgaggaattcattatcacccggcaggaacggagcgggg |
| ChDB3-R | ggtctcgtgggtggtcgccttctgctggtcccgctcgcccgcccaggtc |
| ChDB4-F | gaccagcagaaggcgaccacccacgagacc |
| ChDB4-R | cgtatcacgaggccctttcgtcttc |
| pBesCB-F | taattttgtttaactttaataaggagatataccccatgtccatcactcaagaaacgtttca |
| pBesCB-R | gcagcagcggtttctttaccagactcgagggtacctcacaagggacgcagagcttgct |
| IMDPH-F | tcatcatcatcatcatcacagcagcggccatctagaagtgctttttcagggcccgcatatgctacgtatcgctaaagaagctctgac |
| IMDPH-R | cagcttcctttcgggctttgttagcagccggatcctcaggagcccagacggtagttcggg |
| Prescission-seq-F | ctagaagtgctttttcagggcccg |

#### C. Synthetic gene sequences

WP_030791981 (*Streptomyces lavenduligriseus* BesD)

AGCAGCGGCCATCTAGAAGTGCTTTTTCAGGGCCCGCATATGAGCAGCAATGGGCAGGAAAGCACCGTCGTCAATCC
TCTGGAGCAGGGCGCACTGCGCCGTATGGCGCATCACTACCACCGGTACGGCATCGCCACCGTCACCGATCTGATTC
GGGAAGACGTCCGCAAAAACGTGCGGGCGGAGGCGGACCGGCTGCTGGAGAAATACGCCGAGCGGCGTGATCTCCGC
CTCCAGACCACCGGATACACCCGCCGCTCGATGTCCGTGGTGCAGAGCGAGACGATCGCGGGCAACAGCGAGCTGGT
CACCTCGATCTACGCGAATCCGGAACTGCTCGGCGCGCTGGAGCGCATAGCCGGCGAGAAACTGCACCCCTGCCCCA
AGGCCGACGAGGAATTCCTCATCACCCGGCAGGAACGGAGCGGGGACACGCACGGCTGGCACTGGGGCGATTTCAGC
TTCGCCCTCATCTGGGTGCTCCAGGCCCCGCCCATCGACATCGGCGGCATGCTCCAGTGCGTACCGCACACGGAGTG
GGACAAGTCCGATCCGCGGATCCACCAGTATCTCGTCGACAATCCCATCCACACGTACCACTTCCAGTCGGGCGACG
TGTATTTCCTGCGCACCGACACCACGCTCCACCGCACGGTCCCGCTGCGCGAGGACACCACCCGCATCATCCTGAAC
ATGACCTGGGCGGGCGAGCGGGACCTGCGGCGCGAGCTCAAGGGCGACGACCGCTGGTGGGAGGACGCCGACGTGCC
GGCAGCCGCTGCACTGGATGACTGACTCGAGGATCCGGCTGCTAACAAAGCCCGAAAGGAAGCTGA

SDN46247 (*Streptomyces wuyuanensis* HalB)

AGCAGCGGCCATCTAGAAGTGCTTTTTCAGGGCCCGCATATGACCGAGAACAGCATCACCGCCGCGAATGTCGAAGA
GCTGATCGCGAAGAACATCGCGGAGCGGTTCGCCGACGACCACGAGGTGCTCGGCCTCTCCCAGCACTTCCGCCGCG
AGGGCTACGTCAAGCTCCCCGGCCTGGTCTCCCCGGAGGTCTTCGACGCGGTCGCCGCGGAGACCCACCAGCTGATC
GACACCCACCAGAAGCGCATCGACATCCGTCTCAAGGAGACGGGGGACTCCCCGCGCTACATGTCCACCGTCGGTCA
GAAGGCGATCGCCACCGACGGCTCGCTGATCCCTGCCGTCTACGAGTCCACCGCGCTCAAGGGCTTCCTTTCCCGCC
TCGCCAAGGAGGAGGTCATGGGCTGCCCGTGGGACGAGGAGAAGTACATCATCACCCGCCAGCACCAGAAGGGCGAC
ACCCACGGCTGGCACTGGGGCGACTTCAGCTTCACCGTCATCTGGCTGATCGAGGCCCCGTCGCTGGAGTACGGCGG
CATGCTGCAGTGCATCCCGCACACCGACTGGAACAAGGACGACCCGCGCGTCGAGGACTACCTGCAGAAGCACCCGA
TCCGCAGCTACGGCCACGCCAAGGGCGACCTCTACCTGCTGCGTTCGGACACCACCCTGCACCGCACGGTCCCGCTG

AACGCCGACAGGACCCGCATCATCCTCAACACCTGCTGGGCCAGCCGCGCGGACCAGCAGAAGGCGACCACCCACGA
GACCATGAACGCGATGTTCGACTGATGAGGATCCGGCTGCTAACAAAGCCCGAAAGGAAGCTG

WP_046063366 (*Pseudomonas kilonensis* HalD)
ATGAGCGAACAAACGTCCAGCTTGGTTATTGAAGTAATGGAGCAGCAACTGGCCAAGCATTTCCAGGCTATTTTGCA
AGATGAAAACCGGATGAAGCAGATTCGTAATGAATTTCGCCGAGACGGTTATTTCAACTTCAAGAACTTTTCGTTTC
TACCGAAAAGGATCTTGGAAAATGTCCACGCCGAAGTCCATGCGTTATTGGATGAATATTCGGTGCGTCGCGATGTT
ACTGTTCCGTCCACAGGCAATACCTACCGCAAGATGTACAACGTCAACCAACCGGAGATCGCCGAAGGCGGGACATT
CATCCCTGCGCTCTACCAATCCGAGTCGCTGCGTAAGTTCCTGGGCAACATCGCTGGCGATGATCTGGCGTCTTGCT
GGGAGCAGGAGCAATACCTGGTCACCAAGCTGAGCCACCCGGGCGATACCCACGGCTGGCATTGGGGGGATTACCCG
TACACGATGATCTGGATCATCGAGGCGCCCGAGGACCCGGCGATTGGCGGCGTGCTCCAGTGCGTGCCACACAGCGA
ATGGGACAAGCAGAACCCGCAGATCTGGCAGTACATCCTCAATAACCCGATCAAGTCCTACCACCATCTCAAGGGTG
ATGTGTATTTCCTCAAGTCCGACACCACGTTGCACCACGTCGTGCCGATTCAGCAGGAAACCACTCGGATAATTCTC
AACACATGCTGGGCCAGCGCCCATGACCGGCGAACCGATGTCGCCCACGAAAGCATCGAAGTGATCTGGGACACCAA
GGCCCGGACCACCGAAGAAGCCTGA

**Supplementary Figure 1. Comparison of enzymes that produce 4-Cl-lysine (BesD and HalA), 5-Cl-lysine (HalB), and 4-Cl-ornithine (HalD).** (A) Sequence identity matrix for BesD, HalA, HalB, and HalD. (B) Sequence alignment of BesD (*S. cattleya*, WP_106433083), HalA (*P. fluorescens*, WP_016975823), HalB (*S. wuyuanensis*, WP_093660971), and HalD (*P. kilonensis*, WP_046063366), were performed in MUSCLE [*1*].

A

| % ID | BesD | HalA | HalB | HalD |
|---|---|---|---|---|
| **BesD** | - | 51.8 | 42.4 | 38.6 |
| **HalA** | 51.8 | - | 41.4 | 38.7 |
| **HalB** | 42.4 | 41.4 | - | 40.8 |
| **HalD** | 38.6 | 38.7 | 40.8 | - |

B

```
HalD    MSEQTSSLVIEVMEQQLAKHFQAILQDENRMKQIRNEFRRDGYFNFKNFSFLPKRILENV
HalB    --MTENSITAANVEELIAKNIAERFADDHEVLGLSQHFRREGYVKLPG--LVSPEVFDAV
HalA    -----MNYVLDEARLQ--EHHATNFPES-SVFALRHEFARNGFIKVRN--IVDDDLREKI
BesD    -------MVGSNRQEL--KDVCAPLEKD-DIRRLSQAFHRFGIVTVTE--LIEPHTRKLV
                .         :     : ..  :  : : * * * ...    ::     . :

HalD    HAEVHALLDEYSVRRDVTVPSTGNTYRKMYNVNQPEIAEGGTFIPALYQSESLRKFLGNI
HalB    AAETHQLIDTHQKRIDIRLKETGDSPRYMSTVGQKAIATDGSLIPAVYESTALKGFLSRL
HalA    TREVNSLIDRQLERRDLHLATTDNTPRYMSVVRSEFIAENSTLINTLSKSKGLLETLSQI
BesD    RAEADRLLDQYAERRDLRLATTDYTRRSMSVVPSETIAANSELVTGLYAHRELLAPLEAI
          *.  *:*    * *: :  *. : * *  * .  ** .. ::  :     *   *  :

HalD    AGDDLASC-WEQEQYLVTKLSHPGDTHGWHWGDYPYTMIWIIEAPEDPAIGGVLQCVPHS
HalB    AKEEVMGCPWDEEKYIITRQHQKGDTHGWHWGDFSFTVIWLIEAPSLEY-GGMLQCIPHT
HalA    AGTQLIASVSKDEEYLITKQERKGDTHGWHWGDYSFALIWIIETPSIAK-GGMLQCVPHT
BesD    AGERLHPCPKADEEFLITRQEQRGDTHGWHWGDFSFALIWVLQAPPIDV-GGLLQCVPHT
        *   :  .   :*::::*.  . **********:.:::**::::*     **:***:**:

HalD    EWDKQNPQIWQYILNNPIKSYHHLKGDVYFLKSDTTLHHVVPIQQETTRIILNTCWASAH
HalB    DWNKDDPRVEDYLQKHPIRSYGHAKGDLYLLRSDTTLHRTVPLNADRTRIILNTCWASRA
HalA    SWDKSNPRIHELLCSNPIATYGFVTGDIYFLRTDTTLHRTIPLNEDATRIILNMTWAAEK
BesD    TWDKASPQINRYLVENPIDTYHFESGDVYFLRTDTTLHRTIPLREDTTRIILNMTWAGER
         *:* .*.:   : .:** :* . .**:*:*.:*****..:*:. : ******  **.

HalD    DRRTDVAHESIEVIWDTKARTTEEA---
HalB    DQQKATTHETMNAMFD------------
HalA    DLSRNLHGN--DRWWEDQHVEAAKSLT-
BesD    DLSRKLAAD--DRWWDNAEVSAARAIKD
        *       :  :  ::
```

**Supplementary Table 2.** Data collection and refinement parameters for HalB from *Streptomyces wuyuanensis* (PDB ID 7U6I).

| | HalB, glycine-bound ChainA, apo ChainB |
|---|---|
| **Data collection** | |
| Space group | P $6_2$ 2 2 |
| Cell dimensions | |
| *a*, *b*, *c* (Å) | 165.48, 165.48, 105.33 |
| α, β, γ (°) | 90, 90, 120 |
| Resolution (Å) | 82.74–2.05 (2.123–2.05)* |
| $R_{sym}$ or $R_{merge}$ | 0.16 (3.922) |
| $I / \sigma I$ | 23.9 (1.52) |
| Completeness (%) | 99.8 (97.3) |
| Redundancy | 39.65 (39.3) |
| $CC_{1/2}$ | 1.00 (0.706) |
| **Refinement** | |
| Resolution (Å) | 82.74–2.05 (2.123–2.05)* |
| No. reflections | 2134449 (156810) |
| $R_{work}$ / $R_{free}$ | 0.195/0.221 |
| No. atoms | 4340 |
| Protein | 3879 |
| Ligand/ion | 22 |
| Water | 439 |
| *B*-factors | |
| Protein | 46.12 |
| Ligand/ion | 59.29 |
| Glycine | 52.5 |
| Succinate | 58.6 |
| Water | 56.56 |
| R.m.s. deviations | |
| Bond lengths (Å) | 0.002 |
| Bond angles (°) | 0.53 |

*Values in parentheses are for highest-resolution shell.

**Supplementary Table 3.** Data collection and refinement parameters for HalB from *Streptomyces wuyuanensis* (PDB ID 7U6J).

| | HalB, lysine-bound |
|---|---|
| **Data collection** | |
| Space group | $P12_11$ |
| Cell dimensions | |
| *a*, *b*, *c* (Å) | 70.241, 136.079, 116.24 |
| α, β, γ (°) | 90, 100.016, 90 |
| Resolution (Å) | 69.17–1.9 (1.968–1.9)* |
| $R_{sym}$ or $R_{merge}$ | 0.17 (1.787) |
| $I / \sigma I$ | 8.02 (1.41) |
| Completeness (%) | 99.57 (98.8) |
| Redundancy | 6.9 (6.95) |
| $CC_{1/2}$ | 0.997 (0.497) |
| **Refinement** | |
| Resolution (Å) | 69.17–2.0 (1.968–1.9) |
| No. reflections | 1163368 (115447) |
| $R_{work}$ / $R_{free}$ | 0.1620/0.2114 |
| No. atoms | 18269 |
| Protein | 15867 |
| Ligand/ion | 64 |
| Water | 2338 |
| *B*-factors | |
| Protein | 26.91 |
| Ligand/ion | 28.35 |
| Lysine | 24.43 |
| Succinate | 29 |
| Water | 35.75 |
| R.m.s. deviations | |
| Bond lengths (Å) | 0.006 |
| Bond angles (°) | 0.81 |

*Values in parentheses are for highest-resolution shell.

**Supplementary Figure 2. $Fe^{II}$/αKG-dependent halogenases favor halogenation over hydroxylation.** The chloride ligand binds directly to Fe in halogenases. Upon oxygen binding, halogenases decarboxylate αKG to generate a $Fe^{IV}$-oxo intermediate for abstracting a hydrogen atom from the substrate, yielding a substrate radical. From this intermediate, rebound of either the halogen or the hydroxyl is possible. However, halogenases favor halogen rebound. The related family of hydroxylases instead coordinate the iron with Asp or Glu, leaving no room for chloride. In this case, the hydroxyl group is the only ligand available for rebound with the substrate radical.

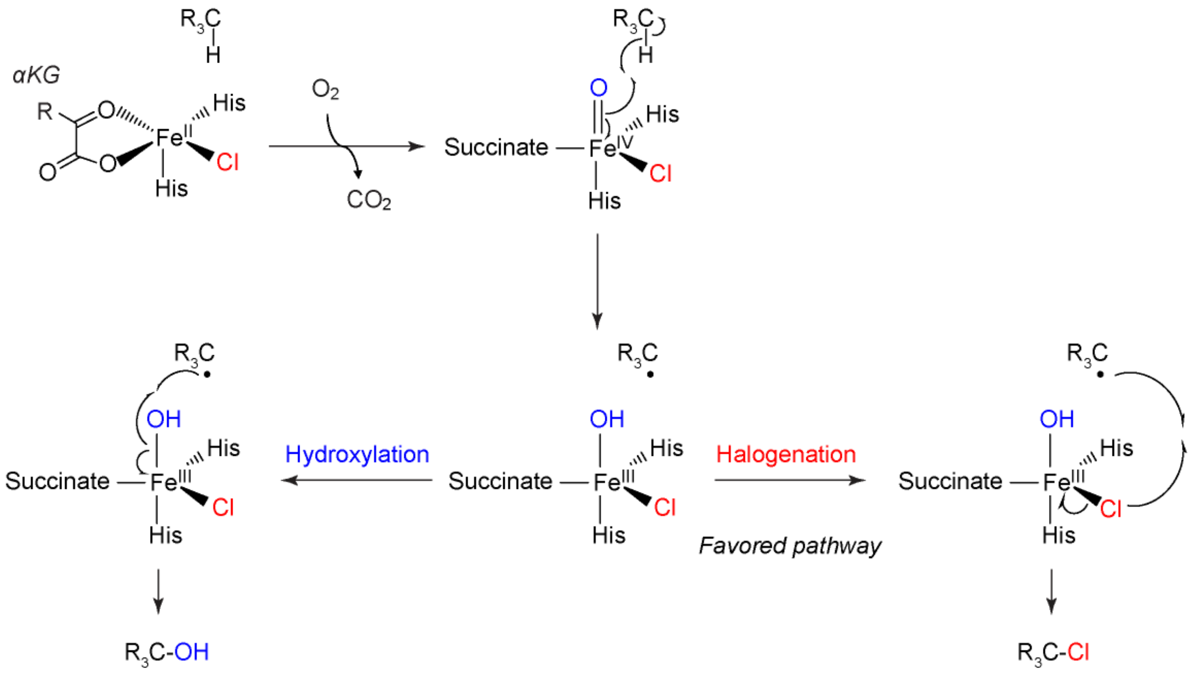

**Supplementary Figure 3.** Propargylglycine (Pra) eventually produced by cells expressing halogenase variants that yield 4-Cl-lysine. Pra reacts with CalFluor 488-azide in the presence of copper (I), ascorbate, and the BTTAA ligand. The resulting propargylglycine-CalFluor product (Pra-CalFluor) is fluorescent ($\lambda_{ex}$ = 485 nm, $\lambda_{em}$ = 528 nm).

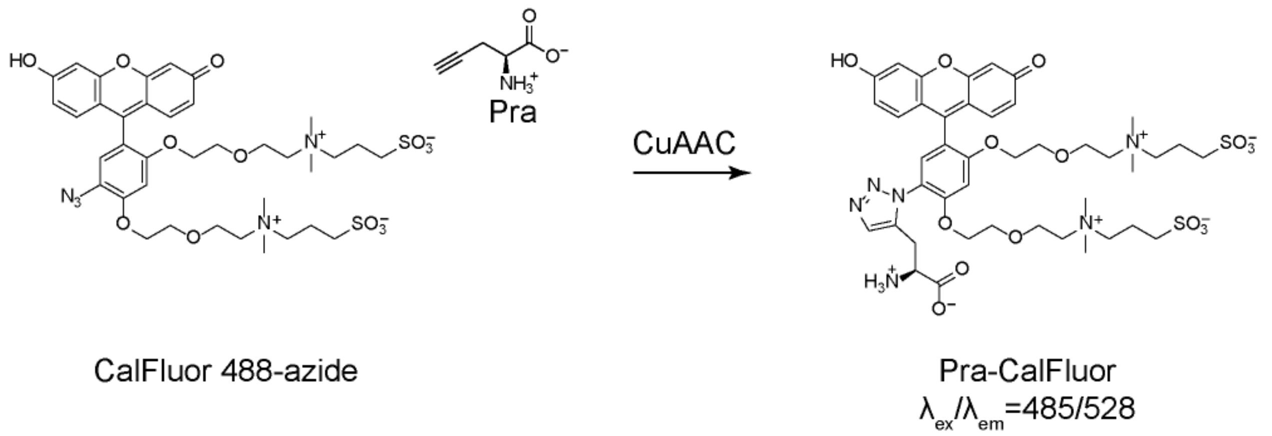

**Supplementary Figure 4. Results of screening the M246NNK library with CuAAC.** Cells expressing BesB, BesC, and a member of the M246NNK library are grown in M9 media supplemented with 0.5 mM lysine. Controls are present on each plate as follows: Lysine hydroxylase (A1-4), HalB (A5-8), BesD (A9-12). After 2 d, cells were supplemented with lysine (0.2 mM), αKG (0.2 mM), and sodium chloride (0.2 mM) and grown for an additional 2 d. Cells were then pelleted by centrifugation and the supernatant was transferred to a fresh plate and mixed 1:1 with CuAAC solution to yield the final reaction mixture (CalFluor 488 azide (1 μM), copper (II) sulfate (0.5 mM), BTTAA (0.1 mM), and sodium ascorbate (5 mM)). Following incubation in the dark for 15 min, the fluorescence was measured using a SynergyMx Microplate Reader (BioTek) at room temperature ($\lambda_{ex}$ = 485 nm, $\lambda_{em}$ = 528 nm). Values for each well are reported relative to the average of wells A1-4 (Hydrox) on each plate.

**Plate 1**

| | 1 | 2 | 3 | 4 | 5 | 6 | 7 | 8 | 9 | 10 | 11 | 12 |
|---|---|---|---|---|---|---|---|---|---|---|---|---|
| **A** | 1.10 | 0.98 | 0.95 | 0.98 | 1.13 | 1.17 | 1.17 | 1.06 | 8.97 | 14.59 | 10.67 | 21.21 |
| **B** | 1.04 | 1.03 | 0.87 | 0.96 | 4.30 | 0.90 | 0.93 | 0.88 | 0.94 | 0.96 | 1.47 | 0.97 |
| **C** | 0.91 | 1.04 | 0.72 | 0.77 | 0.86 | 1.06 | 6.76 | 0.86 | 0.90 | 1.25 | 2.93 | 0.93 |
| **D** | 0.90 | 0.99 | 1.45 | 1.04 | 3.93 | 0.95 | 3.39 | 0.93 | 0.91 | 3.96 | 0.83 | 0.99 |
| **E** | 1.08 | 1.09 | 1.00 | 1.01 | 4.08 | 3.25 | 1.22 | 0.78 | 0.84 | 0.86 | 4.63 | 0.86 |
| **F** | 0.95 | 1.13 | 0.92 | 0.97 | 0.96 | 1.24 | 0.88 | 5.54 | 0.92 | 0.85 | 1.01 | 1.10 |
| **G** | 4.55 | 1.02 | 0.96 | 1.55 | 0.94 | 4.34 | 1.13 | 0.80 | 0.82 | 0.84 | 1.05 | 0.96 |
| **H** | 0.96 | 1.10 | 0.99 | 4.60 | 0.96 | 0.95 | 1.10 | 4.92 | 1.06 | 0.96 | 1.06 | 5.58 |

**Plate 2**

| | 1 | 2 | 3 | 4 | 5 | 6 | 7 | 8 | 9 | 10 | 11 | 12 |
|---|---|---|---|---|---|---|---|---|---|---|---|---|
| **A** | 1.01 | 0.99 | 1.00 | 1.00 | 1.23 | 1.25 | 1.22 | 1.18 | 11.49 | 11.34 | 12.86 | 10.72 |
| **B** | 0.93 | 3.71 | 0.77 | 1.16 | 0.81 | 0.94 | 1.16 | 0.92 | 0.94 | 1.13 | 0.97 | 0.95 |
| **C** | 0.92 | 0.81 | 0.87 | 2.32 | 1.57 | 0.76 | 0.85 | 0.85 | 0.91 | 0.92 | 0.83 | 0.93 |
| **D** | 2.24 | 1.23 | 0.78 | 0.85 | 0.97 | 0.91 | 0.89 | 0.85 | 0.85 | 0.87 | 0.89 | 0.91 |
| **E** | 3.96 | 0.89 | 0.89 | 3.33 | 0.85 | 0.84 | 0.86 | 0.88 | 0.92 | 3.03 | 0.89 | 0.85 |
| **F** | 3.49 | 3.89 | 0.93 | 2.93 | 3.93 | 0.94 | 0.81 | 3.30 | 3.25 | 0.91 | 1.04 | 3.32 |
| **G** | 1.20 | 0.89 | 0.86 | 0.80 | 0.84 | 0.87 | 0.87 | 0.84 | 0.85 | 0.83 | 1.49 | 0.87 |
| **H** | 1.11 | 0.92 | 0.89 | 0.85 | 1.18 | 0.96 | 0.85 | 1.05 | 0.93 | 0.92 | 0.87 | 0.96 |

**Supplementary Table 4. Results of sequencing hits from the M246NNK library screen.** Cells with 3-fold increase in fluorescence relative to the negative control (Hydrox) were grown in LB media. Plasmid DNA was purified from the cells, and sequencing was performed with the Prescission-seq primer, which anneals to the plasmid containing the halogenase variant, but not to the pACYC-BesB-BesC plasmid.

| Plate | Well | M246 | RF |
|---|---|---|---|
| 1 | C11 | T | 2.93 |
| 1 | D10 | T | 3.96 |
| 1 | C7 | V | 6.76 |
| 1 | B5 | L | 4.30 |
| 1 | D7 | T | 3.37 |
| 1 | D5 | L | 3.93 |
| 1 | G1 | L | 4.55 |
| 1 | F8 | V | 5.54 |
| 1 | G6 | L | 4.34 |
| 1 | E5 | T | 4.08 |
| 1 | E6 | T | 3.25 |
| 1 | E11 | T | 4.63 |
| 1 | H12 | T | 5.58 |
| 1 | H8 | I | 4.92 |
| 1 | H4 | T | 4.60 |
| 2 | E1 | I | 3.96 |
| 2 | B2 | L | 3.71 |
| 2 | E4 | V | 3.33 |
| 2 | F2 | L | 3.89 |
| 2 | F12 | I | 3.32 |
| 2 | F8 | L | 3.30 |
| 2 | E10 | L | 3.03 |
| 2 | F9 | L | 3.25 |
| 2 | F5 | I | 3.93 |
| 2 | F4 | I | 2.93 |
| 2 | F1 | L | 3.49 |

**Supplementary Figure 5. Quantification of propargylglycine production by HalB M246 mutants.** (A) A standard curve of CalFluor488 fluorescence in spent M9 media made using commercially available propargylglycine. Media containing Pra was mixed 1:1 with CuAAC solution to yield the final reaction mixture (CalFluor 488 azide (1 μM), copper (II) sulfate (0.5 mM), BTTAA (0.1 mM), and sodium ascorbate (5 mM)). Following incubation in the dark for 15 min, the fluorescence was measured using a SpectraMax M2 Microplate Reader (Molecular Devices) at room temperature ($\lambda_{ex}$ = 485 nm, $\lambda_{em}$ = 528 nm). Data are mean ± sd (n = 3 technical replicates). (B) Quantity of propargylglycine produced by cells containing HalB M246 mutants, BesB, and BesC. Data are mean ± sd (n = 4 technical replicates) Error in [Pra] is obtained by propagation from the error in the curve fit of the standard curve.

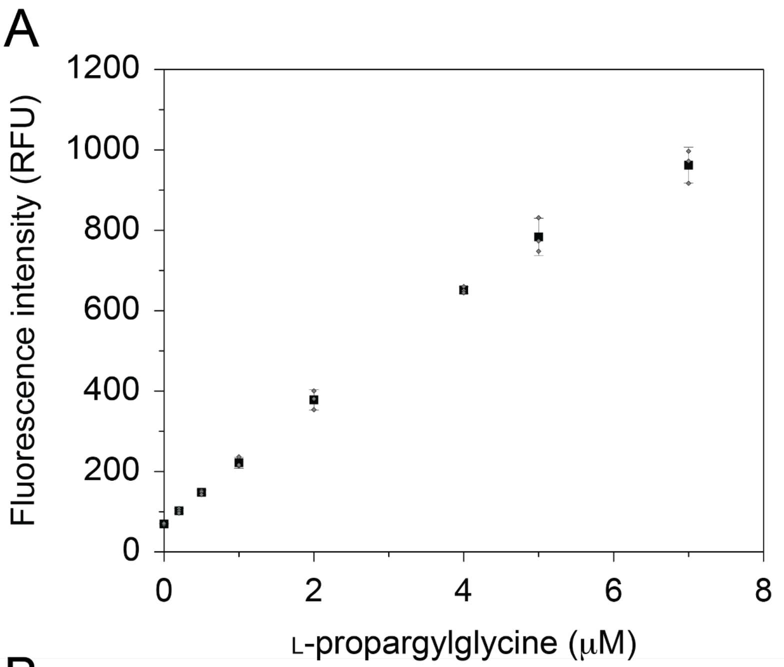

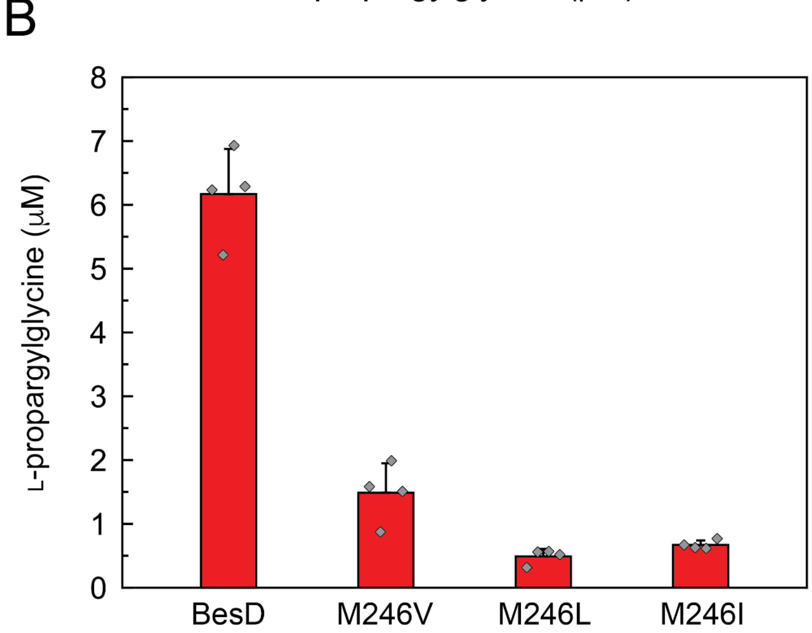

**Supplementary Figure 6. Kinetics of wild type HalB and HalB M246 variants on the substrate lysine.** The rate of succinate formation by HalB variants was monitored by change in $A_{340}$ using an NADH-coupled assay with succinyl-CoA synthetase, pyruvate kinase, and lactate dehydrogenase [*11*]. Reactions were initiated by addition of the halogenase variant (2.5 µM) in the presence of varying concentrations of the lysine substrate (0 - 2 mM). Initial rates of NADH consumption were measured by monitoring $A_{340}$ using a SpectraMax M2 Microplate reader (Molecular Devices) at room temperature. Datapoints are mean ± s.d. (n = 3 technical replicates). Table contains $k_{cat}$, $K_M$, $k_{cat}/K_M$, and $K_i$ calculated by non-linear curve fitting to the Michaelis-Menten equation. $k_{cat}$ and $K_M$ are mean ± s.e. Error in $k_{cat}/K_M$ is obtained by propagation from the individual kinetic terms.

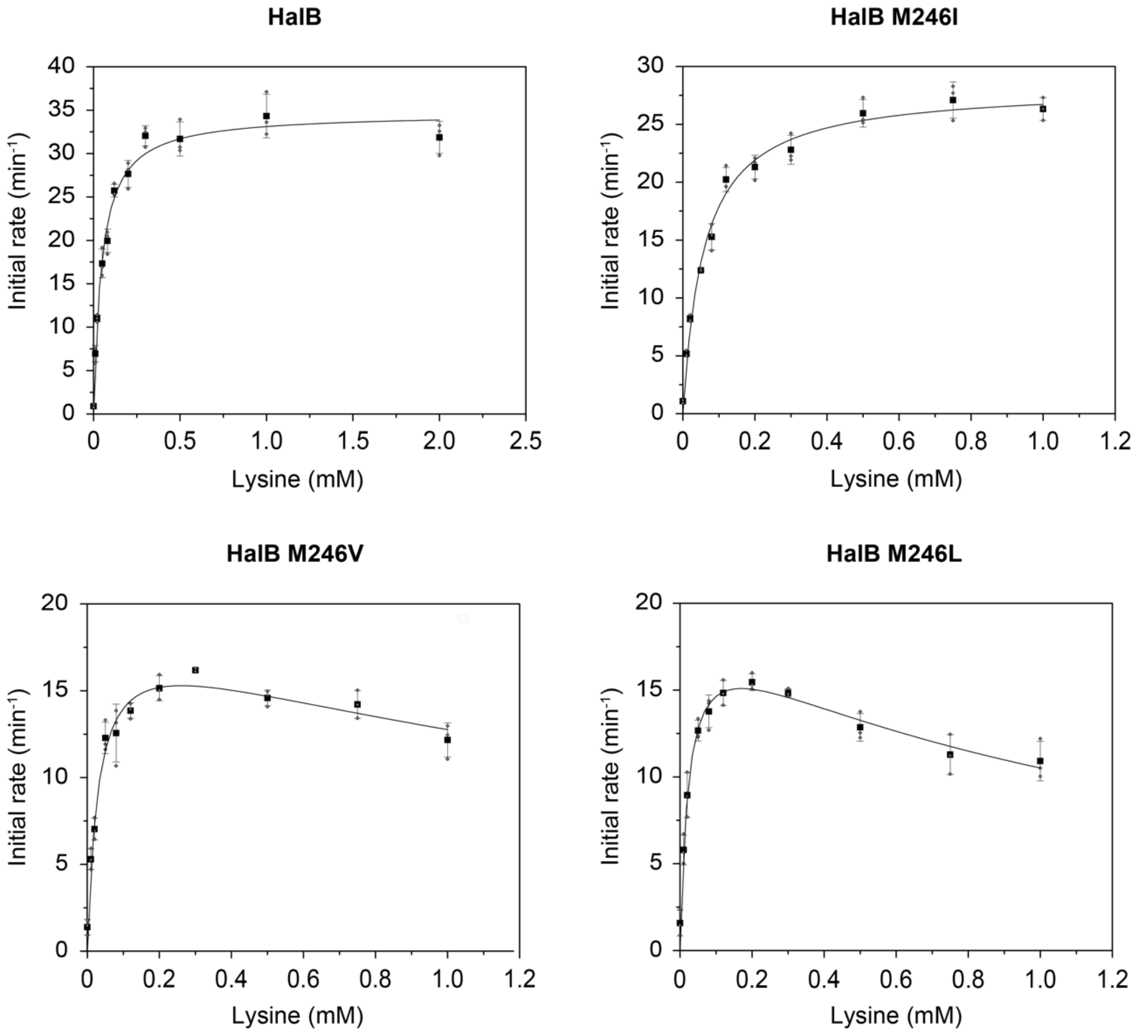

| Enzyme | $k_{cat}$ (min$^{-1}$) | $K_M$ (min$^{-1}$) | $k_{cat}/K_M$ (mM$^{-1}$ min$^{-1}$) | $K_i$ (mM) |
|---|---|---|---|---|
| HalB | 34.7 ± 0.8 | 0.047 ± 0.005 | 741 ± 83 | - |
| HalB M246I | 28.3 ± 0.8 | 0.058 ± 0.007 | 487 ± 59 | - |
| HalB M246L | 19 ± 1 | 0.023 ± 0.004 | 824 ± 140 | 1.2 ± 0.2 |
| HalB M246V | 19 ± 1 | 0.030 ± 0.006 | 620 ± 132 | 2.2 ± 0.7 |

**Supplementary Figure 7. LCMS analysis of 5-Chlorolysine production by HalB variants.** Reactions (50 µL) contained L-lysine · HCl (3 mM), sodium αKG (5 mM), sodium ascorbate (5 mM), $(NH_4)_2Fe(SO_4)_2 \cdot 6H_2O$ (1 mM), and sodium chloride (5 mM) in 100 mM HEPES buffer (pH 7.5). Reactions were initiated by addition of purified BesD or HalB variants (20 µM final concentration) and allowed to proceed for 25 min at room temperature before quenching in 2 vol of methanol with 1% (*v/v*) formic acid. Following centrifugation at 18,000 × *g* for 20 min (4 °C) to remove precipitated protein, samples were analyzed by LC/MS on an Agilent 1290 UPLC-QTOF using the protocol for polar metabolite analysis. 5-Chlorolysine (*m/z* = 181.0738), which was quantified by integrating extracted ion counts at t = 21 min post injection. 4-Cl-lysine elutes at t = 19 min. Due to the instability [*3*], 4-Cl-lysine was quantified by the CuAAC screen in Supplementary Figure 5.

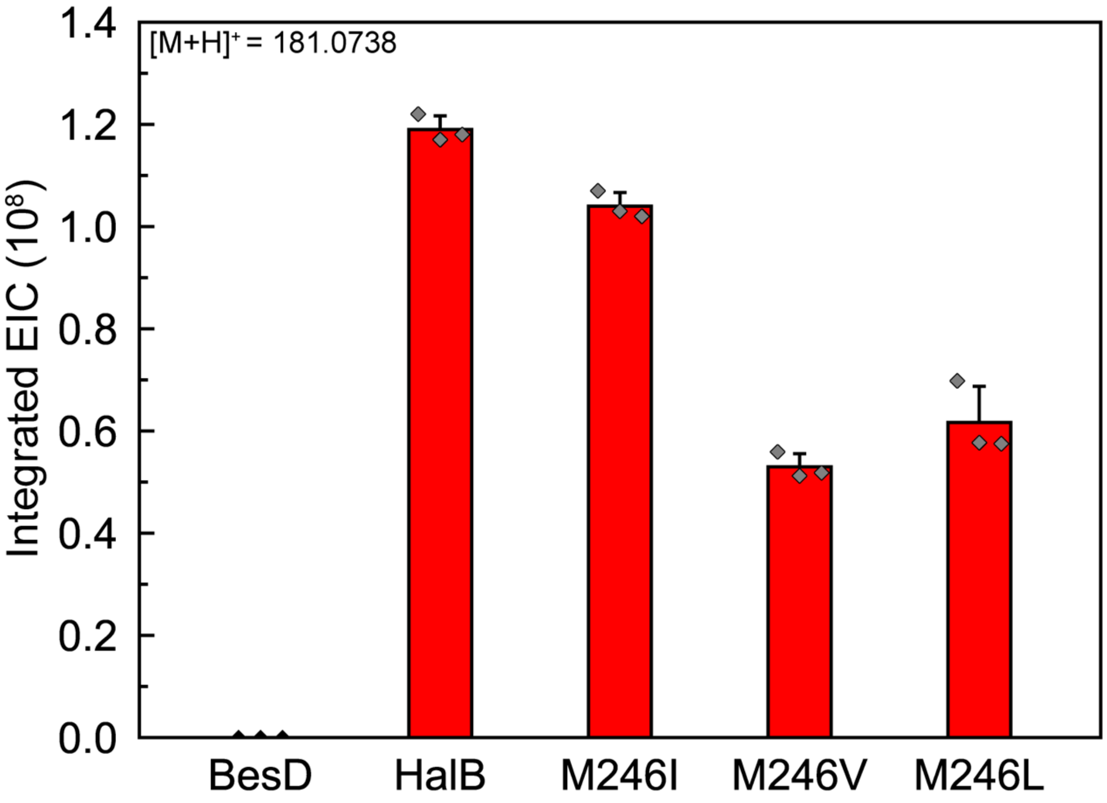

**Supplementary Figure 8. Results of screening the F250NNK libraries with CuAAC.** Cells expressing BesB, BesC, and a member of the F250NNK library are grown in M9 media supplemented with 0.5 mM lysine. Controls are present on each plate as follows: Lysine hydroxylase (A1-4), HalB (A5-8), BesD (A9-12). After 2 d, cells were supplemented with lysine (0.2 mM), αKG (0.2 mM), and sodium chloride (0.2 mM) and grown for an additional 2 d. Cells were then pelleted by centrifugation and the supernatant was transferred to a fresh plate and mixed 1:1 with CuAAC solution to yield the final reaction mixture (CalFluor 488 azide (1 μM), copper (II) sulfate (0.5 mM), BTTAA (0.1 mM), and sodium ascorbate (5 mM)). Following incubation in the dark for 15 min, the fluorescence was measured using a SynergyMx Microplate Reader (BioTek) at room temperature ($\lambda_{ex}$ = 485 nm, $\lambda_{em}$ = 528 nm). Values for each well are reported relative to the average of wells A1-4 (Hydrox) on each plate.

**Plate 1**

| | 1 | 2 | 3 | 4 | 5 | 6 | 7 | 8 | 9 | 10 | 11 | 12 |
|---|---|---|---|---|---|---|---|---|---|---|---|---|
| A | 1.13 | 1.01 | 0.92 | 0.94 | 1.23 | 1.09 | 1.12 | 1.11 | 10.22 | 8.71 | 10.42 | 11.47 |
| B | 1.07 | 1.05 | 0.93 | 0.97 | 0.94 | 1.02 | 0.87 | 1.02 | 0.96 | 0.90 | 0.97 | 0.95 |
| C | 1.21 | 1.23 | 0.79 | 0.82 | 0.70 | 0.73 | 0.80 | 0.82 | 0.79 | 0.56 | 0.88 | 0.55 |
| D | 1.14 | 1.06 | 0.93 | 0.95 | 0.91 | 0.92 | 0.89 | 0.96 | 0.96 | 0.91 | 0.89 | 1.02 |
| E | 1.02 | 0.89 | 0.85 | 1.08 | 0.90 | 0.89 | 0.82 | 0.96 | 0.93 | 0.92 | 0.93 | 1.24 |
| F | 1.21 | 0.87 | 0.90 | 0.88 | 0.86 | 0.89 | 0.82 | 0.88 | 0.95 | 0.91 | 0.92 | 0.93 |
| G | 0.96 | 0.85 | 0.94 | 0.88 | 0.85 | 0.87 | 0.85 | 0.95 | 0.98 | 1.11 | 0.94 | 0.93 |
| H | 0.91 | 1.07 | 0.86 | 0.86 | 0.89 | 0.88 | 0.84 | 0.91 | 0.94 | 1.10 | 0.92 | 0.90 |

**Plate 2**

| | 1 | 2 | 3 | 4 | 5 | 6 | 7 | 8 | 9 | 10 | 11 | 12 |
|---|---|---|---|---|---|---|---|---|---|---|---|---|
| A | 0.97 | 1.04 | 0.99 | 1.00 | 1.03 | 1.08 | 0.97 | 1.05 | 3.53 | 3.81 | 4.58 | 1.96 |
| B | 0.85 | 0.86 | 0.76 | 0.68 | 0.81 | 0.91 | 0.81 | 0.86 | 0.84 | 0.92 | 0.87 | 1.06 |
| C | 1.02 | 0.98 | 0.88 | 0.90 | 0.84 | 0.90 | 0.84 | 0.94 | 0.86 | 0.96 | 0.93 | 0.96 |
| D | 0.90 | 1.13 | 0.88 | 0.87 | 0.86 | 0.86 | 0.78 | 0.91 | 0.93 | 0.92 | 0.95 | 1.19 |
| E | 0.98 | 0.97 | 0.88 | 0.87 | 0.92 | 0.92 | 0.85 | 0.91 | 0.92 | 0.88 | 0.91 | 0.94 |
| F | 1.09 | 0.94 | 0.95 | 1.00 | 0.95 | 0.99 | 0.89 | 0.90 | 0.93 | 0.96 | 0.91 | 0.97 |
| G | 0.92 | 0.90 | 0.94 | 0.97 | 0.87 | 0.95 | 0.84 | 0.93 | 0.99 | 0.99 | 0.78 | 1.00 |
| H | 0.97 | 0.88 | 0.89 | 0.88 | 0.93 | 0.93 | 0.87 | 0.96 | 0.93 | 0.95 | 0.95 | 0.95 |

**Supplementary Figure 9. Results of screening the M246L F250NNK libraries with CuAAC.** Cells expressing BesB, BesC, and a member of the M246L F250NNK library are grown in M9 media supplemented with 0.5 mM lysine. Controls are present on each plate as follows: Lysine hydroxylase (A1-4), HalB (A5-8), BesD (A9-12). After 2 d, cells were supplemented with lysine (0.2 mM), αKG (0.2 mM), and sodium chloride (0.2 mM) and grown for an additional 2 d. Cells were then pelleted by centrifugation and the supernatant was transferred to a fresh plate and mixed 1:1 with CuAAC solution to yield the final reaction mixture (CalFluor 488 azide (1 µM), copper (II) sulfate (0.5 mM), BTTAA (0.1 mM), and sodium ascorbate (5 mM)). Following incubation in the dark for 15 min, the fluorescence was measured using a SynergyMx Microplate Reader (BioTek) at room temperature ($\lambda_{ex}$ = 485 nm, $\lambda_{em}$ = 528 nm). Values for each well are reported relative to the average of wells A1-4 (Hydrox) on each plate.

**Plate 1**

| | 1 | 2 | 3 | 4 | 5 | 6 | 7 | 8 | 9 | 10 | 11 | 12 |
|---|---|---|---|---|---|---|---|---|---|---|---|---|
| A | 1.01 | 1 | 0.99 | 1.01 | 1 | 1.03 | 1 | 1.04 | 7.6 | 3.97 | 9.64 | 6.72 |
| B | 0.94 | 0.85 | 0.94 | 0.73 | 0.68 | 0.68 | 0.67 | 0.62 | 0.76 | 0.78 | 1.25 | 1.14 |
| C | 0.91 | 1.09 | 0.91 | 0.93 | 0.89 | 0.79 | 0.84 | 0.93 | 1.1 | 1.19 | 1.13 | 1.29 |
| D | 0.9 | 0.88 | 1.08 | 0.85 | 0.87 | 0.88 | 0.89 | 1.03 | 0.83 | 1.11 | 0.87 | 0.96 |
| E | 1.33 | 0.85 | 1.11 | 1.07 | 1.18 | 0.9 | 1.06 | 0.82 | 0.91 | 0.89 | 0.88 | 0.93 |
| F | 1 | 1.1 | 0.91 | 1.02 | 0.96 | 0.91 | 1.12 | 0.85 | 1.02 | 1.04 | 0.88 | 0.97 |
| G | 0.92 | 0.92 | 0.91 | 0.91 | 1.27 | 0.91 | 0.91 | 1.05 | 1.18 | 1.51 | 0.93 | 0.96 |
| H | 0.84 | 0.93 | 0.91 | 0.96 | 0.92 | 0.85 | 1.58 | 0.83 | 1.1 | 0.89 | 1.12 | 0.91 |

**Plate 2**

| | 1 | 2 | 3 | 4 | 5 | 6 | 7 | 8 | 9 | 10 | 11 | 12 |
|---|---|---|---|---|---|---|---|---|---|---|---|---|
| A | 0.99 | 1.01 | 1.03 | 0.98 | 1.12 | 1.24 | 1.25 | 1.26 | 7.75 | 7.95 | 7.96 | 8.21 |
| B | 0.89 | 1.02 | 1.08 | 1.69 | 0.83 | 0.91 | 0.93 | 0.92 | 0.99 | 0.96 | 0.81 | 2.94 |
| C | 0.91 | 1.1 | 0.93 | 0.95 | 0.88 | 0.91 | 0.96 | 0.92 | 0.94 | 0.82 | 1.41 | 0.97 |
| D | 0.99 | 0.91 | 0.81 | 0.94 | 0.99 | 0.92 | 1.55 | 2.11 | 1.51 | 0.96 | 0.88 | 2.09 |
| E | 0.9 | 0.9 | 0.95 | 0.88 | 0.93 | 0.83 | 0.88 | 0.89 | 0.92 | 0.89 | 0.84 | 0.93 |
| F | 0.9 | 0.9 | 0.88 | 0.83 | 0.89 | 0.95 | 0.85 | 0.97 | 1.27 | 0.85 | 1.06 | 0.89 |
| G | 1.03 | 0.95 | 0.93 | 0.87 | 0.98 | 0.88 | 0.94 | 1.24 | 2.89 | 0.8 | 1.41 | 0.91 |
| H | 1.93 | 0.96 | 0.83 | 1.02 | 0.92 | 1 | 1.87 | 1.05 | 0.96 | 0.89 | 0.92 | 1.01 |

**Plate 3**

| | 1 | 2 | 3 | 4 | 5 | 6 | 7 | 8 | 9 | 10 | 11 | 12 |
|---|---|---|---|---|---|---|---|---|---|---|---|---|
| A | 1.06 | 1.01 | 0.91 | 1.02 | 1.06 | 1.12 | 1.08 | 1 | 9.6 | 10.86 | 9.89 | 10.34 |
| B | 1.14 | 0.85 | 0.91 | 0.83 | 1.24 | 0.86 | 0.88 | 0.87 | 0.82 | 1.26 | 0.84 | 0.9 |
| C | 0.96 | 1.36 | 1.36 | 1.27 | 0.84 | 0.8 | 0.85 | 0.8 | 0.68 | 1.28 | 0.78 | 0.89 |
| D | 0.89 | 0.87 | 0.72 | 0.76 | 1.24 | 0.67 | 0.81 | 0.77 | 0.79 | 0.76 | 0.72 | 1.12 |
| E | 0.93 | 0.88 | 0.9 | 0.91 | 2.73 | 1.06 | 0.83 | 1.46 | 1.29 | 0.97 | 1.26 | 1.18 |
| F | 0.94 | 0.97 | 1.33 | 0.93 | 0.89 | 1.39 | 3.01 | 0.91 | 0.85 | 1.72 | 0.87 | 1.02 |
| G | 1.96 | 0.96 | 0.86 | 0.93 | 0.96 | 1.02 | 0.93 | 0.92 | 0.85 | 0.94 | 1.42 | 0.91 |
| H | 1.03 | 0.98 | 1.96 | 2.78 | 0.95 | 0.89 | 0.89 | 0.82 | 0.89 | 0.96 | 0.92 | 1.51 |

**Supplementary Table 5. Results of sequencing hits from the M246L F250NNK library screen.** Cells with 2-fold increase in fluorescence relative to the negative control (Hydrox) were grown in LB media. Plasmid DNA was purified from the cells, and sequencing was performed with the Prescission-seq primer, which anneals to the plasmid containing the halogenase variant, but not to the pACYC-BesB-BesC plasmid.

| Plate | Well | M246 | F250 | RF |
|---|---|---|---|---|
| 2 | B12 | L | F | 2.94 |
| 2 | D12 | L | C | 2.09 |
| 2 | D8 | L | C | 2.11 |
| 2 | G9 | L | F | 2.89 |
| 3 | H4 | L | F | 2.78 |
| 3 | E5 | L | F | 2.73 |
| 3 | F7 | L | F | 3.01 |

**Supplementary Figure 10. Results of screening the M246V F250NNK libraries with CuAAC.** Cells expressing BesB, BesC, and a member of the M246V F250NNK library are grown in M9 media supplemented with 0.5 mM lysine. Controls are present on each plate as follows: Lysine hydroxylase (A1-4), HalB (A5-8), BesD (A9-12). After 2 d, cells were supplemented with lysine (0.2 mM), αKG (0.2 mM), and sodium chloride (0.2 mM) and grown for an additional 2 d. Cells were then pelleted by centrifugation and the supernatant was transferred to a fresh plate and mixed 1:1 with CuAAC solution to yield the final reaction mixture (CalFluor 488 azide (1 μM), copper (II) sulfate (0.5 mM), BTTAA (0.1 mM), and sodium ascorbate (5 mM)). Following incubation in the dark for 15 min, the fluorescence was measured using a SynergyMx Microplate Reader (BioTek) at room temperature ($\lambda_{ex}$ = 485 nm, $\lambda_{em}$ = 528 nm). Values for each well are reported relative to the average of wells A1-4 (Hydrox) on each plate.

**Plate 1**

| | 1 | 2 | 3 | 4 | 5 | 6 | 7 | 8 | 9 | 10 | 11 | 12 |
|---|---|---|---|---|---|---|---|---|---|---|---|---|
| **A** | 0.96 | 1.05 | 0.96 | 1.02 | 1.16 | 1.11 | 0.96 | 0.98 | 4.89 | 6.33 | 7.24 | 5.75 |
| **B** | 6.15 | 0.98 | 0.92 | 0.96 | 0.96 | 1.66 | 3.33 | 0.86 | 0.87 | 1.59 | 0.82 | 0.95 |
| **C** | 1.05 | 0.94 | 1.03 | 0.90 | 0.96 | 1.31 | 1.42 | 0.95 | 0.88 | 1.55 | 1.42 | 0.96 |
| **D** | 1.02 | 0.96 | 1.02 | 0.98 | 1.07 | 1.36 | 0.87 | 0.93 | 0.88 | 0.92 | 0.90 | 1.01 |
| **E** | 1.03 | 1.01 | 1.04 | 0.89 | 0.97 | 1.14 | 1.24 | 1.30 | 0.91 | 0.74 | 0.97 | 0.91 |
| **F** | 1.45 | 2.53 | 0.93 | 3.54 | 1.14 | 0.90 | 0.87 | 1.11 | 1.20 | 1.26 | 1.39 | 1.42 |
| **G** | 0.96 | 2.46 | 0.65 | 0.61 | 0.60 | 0.60 | 0.83 | 0.71 | 0.60 | 0.66 | 0.71 | 0.43 |
| **H** | 1.06 | 0.90 | 0.91 | 1.11 | 1.20 | 0.99 | 0.91 | 0.89 | 0.90 | 2.51 | 0.91 | 1.64 |

**Plate 2**

| | 1 | 2 | 3 | 4 | 5 | 6 | 7 | 8 | 9 | 10 | 11 | 12 |
|---|---|---|---|---|---|---|---|---|---|---|---|---|
| **A** | 1.06 | 1.03 | 0.95 | 0.95 | 0.97 | 1.13 | 1.01 | 1.20 | 9.77 | 9.75 | 11.61 | 10.56 |
| **B** | 1.07 | 0.97 | 2.36 | 0.97 | 0.98 | 1.63 | 0.94 | 4.03 | 0.88 | 1.03 | 1.17 | 0.91 |
| **C** | 1.07 | 0.98 | 0.89 | 0.92 | 0.94 | 2.26 | 0.93 | 0.87 | 0.74 | 0.82 | 1.38 | 1.11 |
| **D** | 0.89 | 1.30 | 0.82 | 1.39 | 0.71 | 0.81 | 0.82 | 0.87 | 1.19 | 1.00 | 0.92 | 0.92 |
| **E** | 2.32 | 0.98 | 1.64 | 2.23 | 0.87 | 0.80 | 0.78 | 0.73 | 2.01 | 0.79 | 1.45 | 0.90 |
| **F** | 1.36 | 1.29 | 0.91 | 0.85 | 0.87 | 0.92 | 1.18 | 1.41 | 1.57 | 0.91 | 1.07 | 1.06 |
| **G** | 0.91 | 0.85 | 1.00 | 0.87 | 0.78 | 1.01 | 1.72 | 1.11 | 0.99 | 0.85 | 1.10 | 0.73 |
| **H** | 1.10 | 1.01 | 2.29 | 0.94 | 0.98 | 0.96 | 0.90 | 1.05 | 0.92 | 0.99 | 1.14 | 1.63 |

**Supplementary Table 6. Results of sequencing hits from the M246V F250NNK library screen.** Cells with 2-fold increase in fluorescence relative to the negative control (Hydrox) were grown in LB media. Plasmid DNA was purified from the cells, and sequencing was performed with the Prescission-seq primer, which anneals to the plasmid containing the halogenase variant, but not to the pACYC-BesB-BesC plasmid.

| Plate | Well | M246 | F250 | RF |
|---|---|---|---|---|
| 1 | B1 | V | F | 6.15 |
| 1 | F2 | V | C | 2.53 |
| 1 | G2 | V | T | 2.46 |
| 1 | F4 | V | F | 3.54 |
| 1 | B7 | V | T | 3.33 |
| 1 | H10 | V | F | 2.51 |
| 2 | E1 | V | T | 2.32 |
| 2 | B3 | V | T | 2.36 |
| 2 | H3 | V | T | 2.29 |
| 2 | E4 | V | T | 2.23 |
| 2 | C6 | V | T | 2.26 |
| 2 | B8 | V | F | 4.03 |

**Supplementary Figure 11. Results of screening the M246I F250NNK libraries with CuAAC.** Cells expressing BesB, BesC, and a member of the M246I F250NNK library are grown in M9 media supplemented with 0.5 mM lysine. Controls are present on each plate as follows: Lysine hydroxylase (A1-4), HalB (A5-8), BesD (A9-12). After 2 d, cells were supplemented with lysine (0.2 mM), αKG (0.2 mM), and sodium chloride (0.2 mM) and grown for an additional 2 d. Cells were then pelleted by centrifugation and the supernatant was transferred to a fresh plate and mixed 1:1 with CuAAC solution to yield the final reaction mixture (CalFluor 488 azide (1 µM), copper (II) sulfate (0.5 mM), BTTAA (0.1 mM), and sodium ascorbate (5 mM)). Following incubation in the dark for 15 min, the fluorescence was measured using a SynergyMx Microplate Reader (BioTek) at room temperature ($\lambda_{ex}$ = 485 nm, $\lambda_{em}$ = 528 nm). Values for each well are reported relative to the average of wells A1-4 (Hydrox) on each plate.

**Plate 1**

| | 1 | 2 | 3 | 4 | 5 | 6 | 7 | 8 | 9 | 10 | 11 | 12 |
|---|---|---|---|---|---|---|---|---|---|---|---|---|
| **A** | 1.01 | 0.99 | 1.00 | 1.01 | 1.09 | 1.10 | 1.08 | 1.09 | 12.01 | 6.45 | 11.64 | 8.23 |
| **B** | 0.86 | 1.24 | 0.94 | 0.86 | 0.92 | 0.94 | 0.99 | 0.86 | 0.85 | 0.82 | 1.47 | 1.00 |
| **C** | 0.88 | 0.82 | 0.88 | 0.98 | 0.86 | 0.83 | 1.26 | 0.87 | 1.16 | 0.89 | 1.22 | 0.87 |
| **D** | 0.94 | 1.37 | 1.48 | 1.54 | 0.79 | 1.39 | 1.83 | 0.90 | 0.77 | 1.01 | 1.21 | 0.82 |
| **E** | 0.80 | 0.89 | 0.87 | 0.85 | 0.86 | 0.94 | 1.05 | 0.87 | 0.88 | 0.79 | 0.86 | 3.21 |
| **F** | 1.06 | 0.75 | 0.97 | 1.04 | 0.82 | 1.55 | 1.01 | 2.54 | 0.86 | 1.29 | 0.89 | 1.43 |
| **G** | 0.85 | 0.82 | 0.86 | 0.89 | 1.03 | 0.90 | 0.85 | 0.85 | 3.51 | 0.88 | 0.93 | 0.95 |
| **H** | 0.78 | 0.83 | 1.70 | 0.90 | 0.89 | 0.84 | 0.86 | 0.94 | 0.92 | 0.92 | 0.90 | 0.93 |

**Plate 2**

| | 1 | 2 | 3 | 4 | 5 | 6 | 7 | 8 | 9 | 10 | 11 | 12 |
|---|---|---|---|---|---|---|---|---|---|---|---|---|
| **A** | 0.95 | 0.99 | 1.01 | 1.04 | 1.08 | 1.04 | 1.05 | 1.01 | 3.72 | 1.76 | 3.50 | 3.97 |
| **B** | 0.93 | 0.87 | 0.90 | 0.90 | 0.86 | 0.90 | 0.85 | 1.00 | 1.59 | 0.85 | 0.90 | 1.74 |
| **C** | 1.06 | 0.88 | 0.85 | 0.96 | 0.92 | 0.92 | 0.88 | 0.91 | 0.91 | 0.88 | 0.87 | 0.97 |
| **D** | 0.92 | 0.86 | 0.85 | 1.14 | 0.87 | 0.95 | 0.91 | 1.01 | 0.88 | 0.86 | 0.91 | 0.89 |
| **E** | 0.79 | 0.89 | 0.89 | 0.82 | 1.08 | 0.92 | 0.91 | 0.94 | 0.92 | 1.02 | 0.89 | 0.90 |
| **F** | 0.86 | 0.90 | 0.92 | 0.91 | 1.00 | 0.97 | 0.91 | 0.89 | 0.93 | 0.97 | 0.92 | 1.14 |
| **G** | 0.91 | 0.87 | 0.94 | 0.95 | 1.08 | 0.92 | 0.97 | 0.93 | 0.90 | 1.00 | 0.90 | 0.89 |
| **H** | 0.84 | 0.91 | 0.86 | 0.85 | 1.12 | 0.95 | 0.86 | 0.98 | 0.93 | 0.90 | 0.94 | 1.18 |

**Supplementary Table 7. Results of sequencing hits from the M246I F250NNK library screen.** Cells with 2-fold increase in fluorescence relative to the negative control (Hydrox) were grown in LB media. Plasmid DNA was purified from the cells, and sequencing was performed with the Prescission-seq primer, which anneals to the plasmid containing the halogenase variant, but not to the pACYC-BesB-BesC plasmid.

| Plate | Well | M246 | F250 | RF |
|---|---|---|---|---|
| 1 | F8 | I | F | 2.54 |
| 1 | G9 | I | F | 3.51 |
| 1 | E12 | I | F | 3.21 |
| 2 | B9 | I | F | 1.59 |
| 2 | B12 | I | F | 1.74 |

**Supplementary Table 8.** Data collection and refinement parameters for HalD from *Pseudomonas kilonensis.* (PDB ID 7U6H).

| | HalD, ornithine-bound |
|---|---|
| **Data collection** | |
| Space group | P1 |
| Cell dimensions | |
| *a*, *b*, *c* (Å) | 72.0505, 73.226, 73.176 |
| α, β, γ (°) | 66.2385, 75.7995, 85.279 |
| Resolution (Å) | 69.84–2.0 (2.071–2.0)* |
| $R_{sym}$ or $R_{merge}$ | 0.1333 (2.019) |
| $I / \sigma I$ | 8.84 (1.49) |
| Completeness (%) | 97.46 (96.02) |
| Redundancy | 7.1 (6.7) |
| $CC_{1/2}$ | 0.998 (0.541) |
| **Refinement** | |
| Resolution (Å) | 69.84-2.0 |
| No. reflections | 620918 (57756) |
| $R_{work}$ / $R_{free}$ | 0.18/0.21 |
| No. atoms | 9369 |
| Protein | 8434 |
| Ligand/ion | 159 |
| Water | 776 |
| *B*-factors | |
| Protein | 44.09 |
| Ligand/ion | 64.17 |
| Metal | 33.75 |
| Cl | 43.5 |
| Ornithine | 40.19 |
| αKG | 45.69 |
| Water | 50.98 |
| R.m.s. deviations | |
| Bond lengths (Å) | 0.005 |
| Bond angles (°) | 0.98 |

*Values in parentheses are for highest-resolution shell.

**Supplementary Figure 12. Predicted stereochemistry of hydrogen atom abstraction.** In the crystal structures of BesD (A), HalB (B), and HalD (C), the pro-R hydrogen atom of the substrate carbon is positioned towards the metal and halide binding sites. Following hydrogen atom abstraction, it is anticipated that barring substrate rearrangement, the resulting radical rebound would favor halogenation with R stereochemistry. This outcome has been verified by NMR in the case of BesD [*12*]. For HalB (B), the αKG, Fe, and Cl have been included by alignment with BesD for clarity.

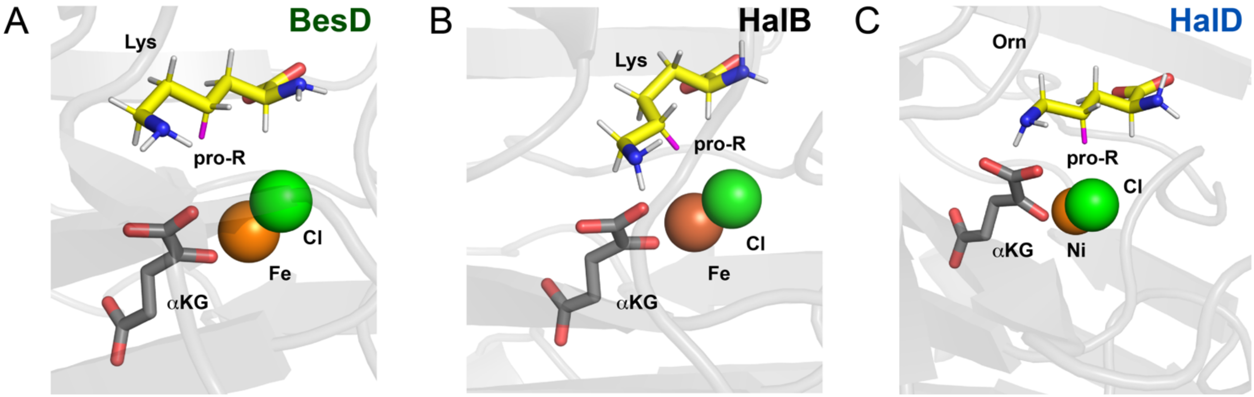

**Supplementary Figure 13. Comparison of BesD, HalB, and HalD.** Overall fold comparison of BesD (green), HalB (grey), and HalD (blue) with substrate covering lids shown in magenta. The substrate covering lid of HalD is more structurally similar to the lid in HalB than to the lid in BesD.

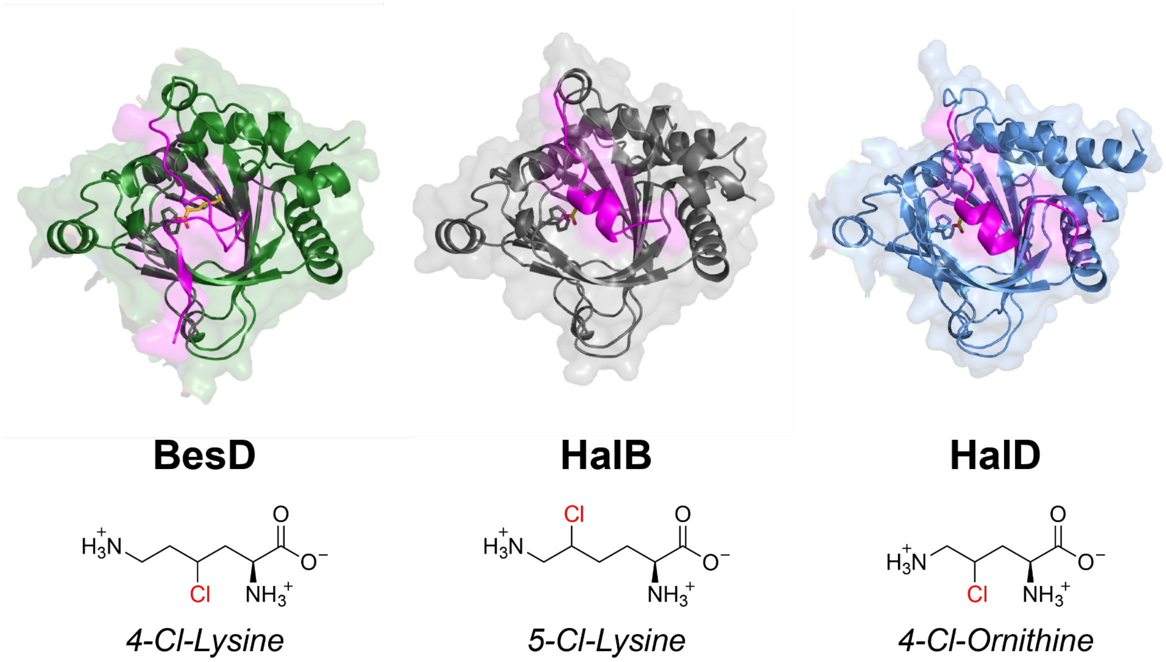

**Supplementary Figure 14. Active site comparison of HalD (blue) and BesD (green).** (A) An overlay of the active sites of HalD and BesD reveals high similarity in the carboxylate-binding and ε-amine-binding residues. In both cases, the substrate is modified at $C_4$. However, the structure of the substrate binding pocket that lines the aliphatic sidechain differs between the two enzymes. (B) In the ornithine halogenase, HalD, the substrate binding pocket is composed of Ile250, Ile253, and Trp254. (C) In the lysine 4-halogenase, BesD, the substrate binding pocket is composed of Trp238 and Trp239.

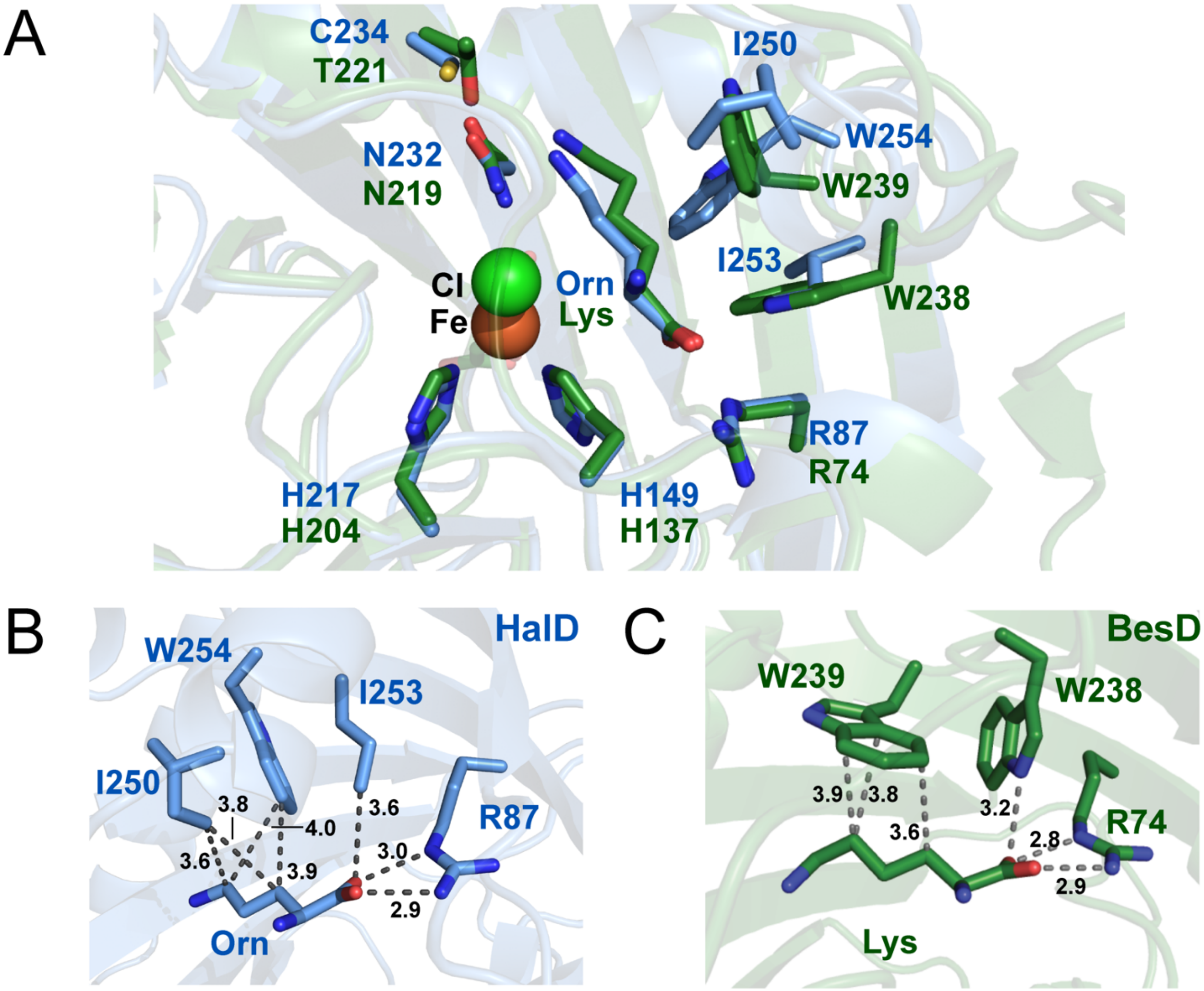

**Supplementary Figure 15. Purification of halogenase variants.** Lane **1**, HalB; **2**, HalD; **3**, HalB F250W; **4**, HalD W254F; **5**, HalB M246V; **6**, HalB M246L; **7**, HalB M246I; **8**, HalB M246I M249I F250W; **9**, HalD I250M I253M W254F. The purification was performed once.

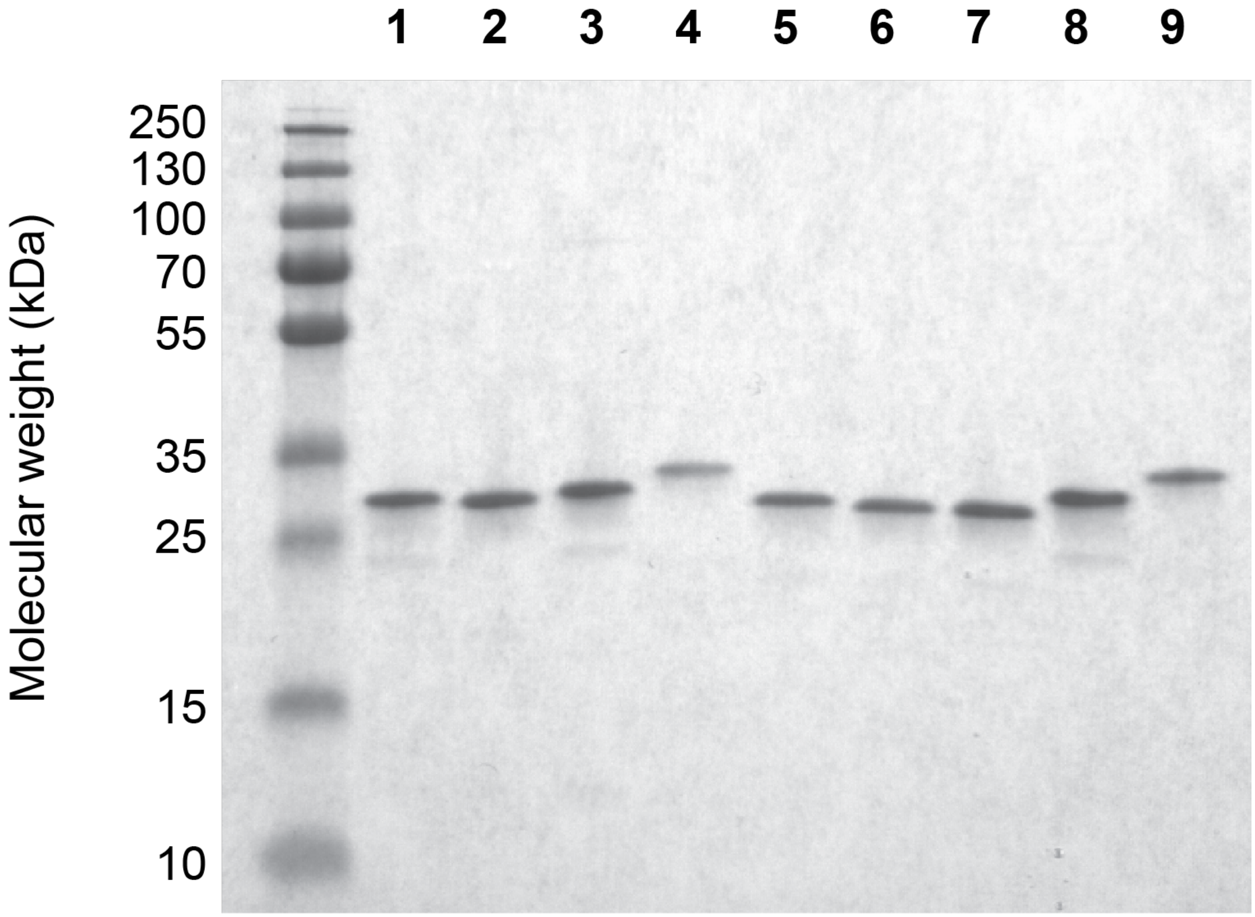

**Supplementary Figure 16. Kinetics of wild type HalB on lysine and ornithine.** The rate of succinate formation by HalB was monitored by change in $A_{340}$ using an NADH-coupled assay with succinyl-CoA synthetase, pyruvate kinase, and lactate dehydrogenase [*11*]. Reactions were initiated by addition of HalB (2.5 μM) in the presence of varying concentrations of the lysine (0 - 2 mM) or ornithine (0 - 0.8 mM). Initial rates of NADH consumption were measured by monitoring $A_{340}$ using a SpectraMax M2 Microplate reader (Molecular Devices) at room temperature. Datapoints are mean ± s.d. (n = 3 technical replicates). Table contains $k_{cat}$, $K_M$, $k_{cat}/K_M$, and $K_i$ calculated by non-linear curve fitting to the Michaelis-Menten equation. $k_{cat}$ and $K_M$ are mean ± s.e. Error in $k_{cat}/K_M$ is obtained by propagation from the individual kinetic terms.

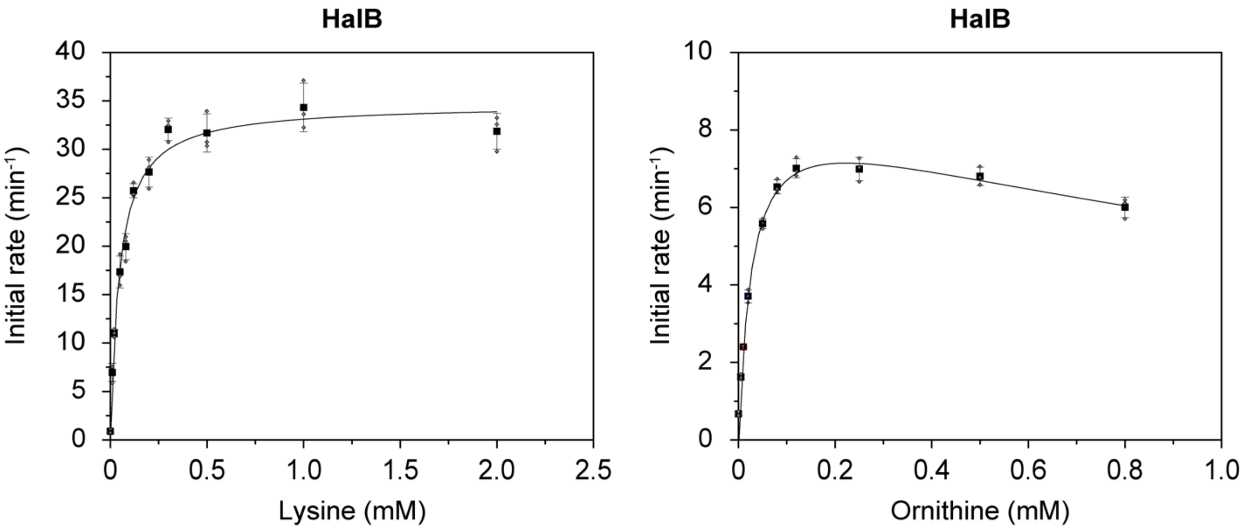

| Enzyme | Substrate | $k_{cat}$ ($min^{-1}$) | $K_M$ ($min^{-1}$) | $k_{cat}/K_M$ ($mM^{-1}$ $min^{-1}$) | $K_i$ (mM) |
|---|---|---|---|---|---|
| HalB | Lysine | 34.7 ± 0.8 | 0.047 ± 0.005 | 741 ± 83 | - |
| HalB | Ornithine | 8.9 ± 0.5 | 0.026 ± 0.004 | 335 ± 54 | 1.8 ± 0.5 |

**Supplementary Figure 17. Kinetics of HalD W254F on lysine and ornithine.** The rate of succinate formation by HalB was monitored by change in $A_{340}$ using an NADH-coupled assay with succinyl-CoA synthetase, pyruvate kinase, and lactate dehydrogenase [*11*]. Reactions were initiated by addition of HalB W254F (2.5 µM) in the presence of varying concentrations of the lysine (0-5 mM) or ornithine (0 - 2 mM). Initial rates of NADH consumption were measured by monitoring $A_{340}$ using a SpectraMax M2 Microplate reader (Molecular Devices) at room temperature. Datapoints are mean ± s.d. (n = 3 technical replicates). Table contains $k_{cat}$, $K_M$, $k_{cat}/K_M$, and $K_i$ calculated by non-linear curve fitting to the Michaelis-Menten equation. $k_{cat}$ and $K_M$ are mean ± s.e. Error in $k_{cat}/K_M$ is obtained by propagation from the individual kinetic terms.

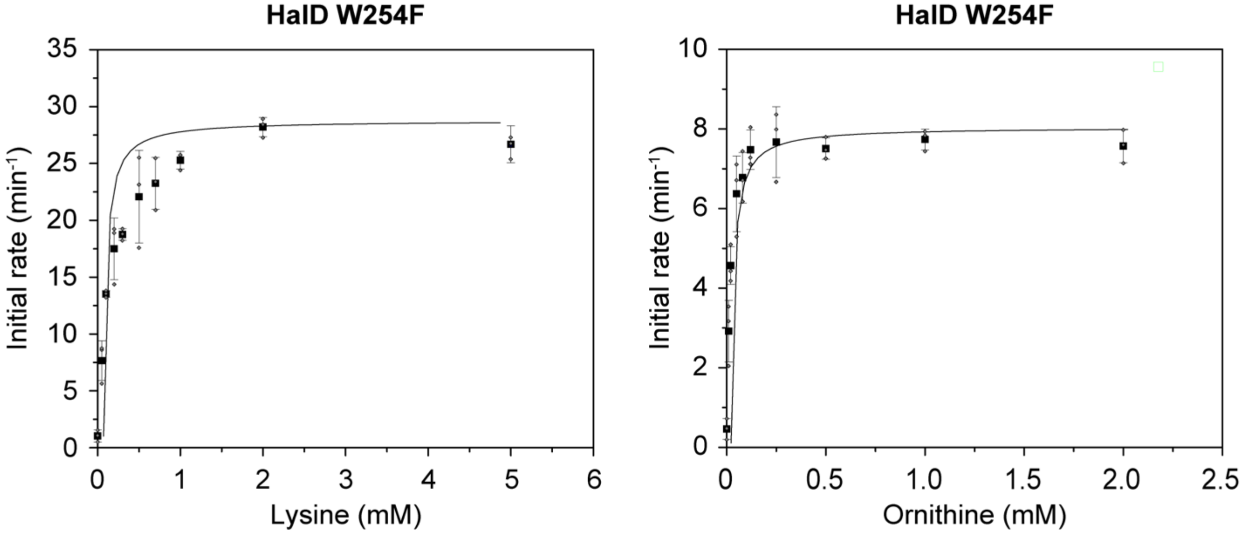

| Enzyme | Substrate | $k_{cat}$ (min$^{-1}$) | $K_M$ (min$^{-1}$) | $k_{cat}/K_M$ (mM$^{-1}$ min$^{-1}$) | $K_i$ (mM) |
|---|---|---|---|---|---|
| HalB | Lysine | 34.7 ± 0.8 | 0.047 ± 0.005 | 741 ± 83 | - |
| HalB | Ornithine | 8.9 ± 0.5 | 0.026 ± 0.004 | 335 ± 54 | 1.8 ± 0.5 |
| HalD | Lysine | 16.16 ± 0.43 | 1.22 ± 0.12 | 13.24 ± 3.6 | - |
| HalD | Ornithine | 10.14 ± 0.28 | 0.03 ± 0.004 | 327.1 ± 70 | - |
| HalD W254F | Lysine | 28.2 ± 0.7 | 0.13 ± 0.01 | 222 ± 24 | - |
| HalD W254F | Ornithine | 7.9 ± 0.1 | 0.015 ± 0.002 | 547 ± 63 | - |

**Supplementary Figure 18. Kinetics of HalD I250M I253M W254F on lysine and ornithine.** The rate of succinate formation by HalB was monitored by change in $A_{340}$ using an NADH-coupled assay with succinyl-CoA synthetase, pyruvate kinase, and lactate dehydrogenase [*11*]. Reactions were initiated by addition of HalB W254F (2.5 μM) in the presence of varying concentrations of the lysine (0-5 mM) or ornithine (0-2 mM). Initial rates of NADH consumption were measured by monitoring $A_{340}$ using a SpectraMax M2 Microplate reader (Molecular Devices) at room temperature. Datapoints are mean ± s.d. (n = 3 technical replicates). Table contains $k_{cat}$, $K_M$, $k_{cat}/K_M$, and $K_i$ calculated by non-linear curve fitting to the Michaelis-Menten equation. $k_{cat}$ and $K_M$ are mean ± s.e. Error in $k_{cat}/K_M$ is obtained by propagation from the individual kinetic terms.

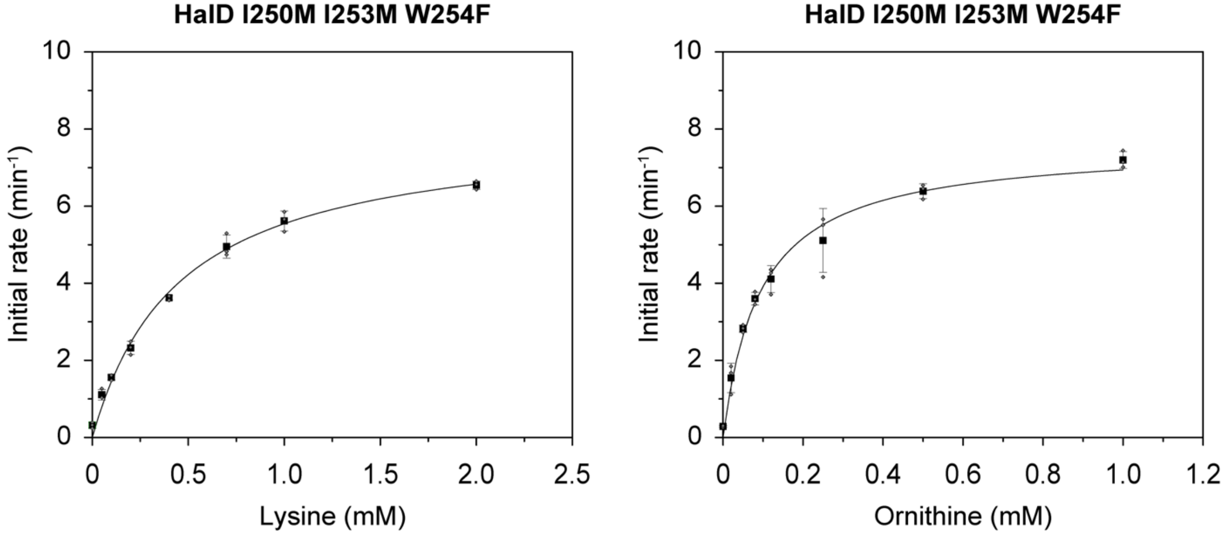

| Enzyme | Substrate | $k_{cat}$ (min$^{-1}$) | $K_M$ (min$^{-1}$) | $k_{cat}/K_M$ (mM$^{-1}$ min$^{-1}$) | $K_i$ (mM) |
|---|---|---|---|---|---|
| HalB | Lysine | 34.7 ± 0.8 | 0.047 ± 0.005 | 741 ± 83 | - |
| HalB | Ornithine | 8.9 ± 0.5 | 0.026 ± 0.004 | 335 ± 54 | 1.8 ± 0.5 |
| HalD | Lysine | 16.16 ± 0.43 | 1.22 ± 0.12 | 13.24 ± 3.6 | - |
| HalD | Ornithine | 10.14 ± 0.28 | 0.03 ± 0.004 | 327.1 ± 70 | - |
| HalD W254F | Lysine | 28.2 ± 0.7 | 0.13 ± 0.01 | 222 ± 24 | - |
| HalD W254F | Ornithine | 7.9 ± 0.1 | 0.015 ± 0.002 | 547 ± 63 | - |
| HalD I250M I253M W254F | Lysine | 8.1 ± 0.4 | 0.45 ± 0.05 | 18 ± 2 | - |
| HalD I250M I253M W254F | Ornithine | 7.6 ± 0.3 | 0.09 ± 0.01 | 81 ± 10 | - |
